# Genome-wide classification of tumor-derived reads from bulk long-read sequencing

**DOI:** 10.64898/2026.03.03.709085

**Authors:** Toby M. Baker, Nedas Matulionis, Cassidy Andrasz, Dan Gerke, Natalia Garcia-Dutton, David Atkinson, Kami Chiotti, Selina Wu, Suchita Lulla, Jieun Oh, Helena Winata, Rong-Rong Huang, Jenny Lester, Beth Y. Karlan, Paul T. Spellman

## Abstract

DNA extracted from tissue samples typically derives from a complex mixture of cell types. Without single cell analysis, it has been generally impossible to determine the cell type of origin for most molecules. One clear example of this is in the complex milieu of a human neoplasm. Here, we develop ROCIT (https://github.com/tobybaker/rocit), a transformer-based model to classify the tumor or non-tumor origin of individual reads from bulk tumor samples sequenced with long-read whole genome sequencing. Using somatic mutations to derive training data, ROCIT uses read-level methylation patterns to accurately classify reads from anywhere in the genome without requiring the adjacent normal tissue or the explicit identification of tumor differentially methylated regions. We apply ROCIT to a cohort of prostate and ovarian tumors and demonstrate high classification accuracy across the entire genome. We then demonstrate the potential of ROCIT predictions to improve somatic variant calling. ROCIT represents a major step forward in the analysis of bulk tumors with long-reads, enabling the accurate and sensitive identification of reads with specific cell types of origin genome-wide.

## Introduction

It is frequently useful to distinguish the cell of origin of individual DNA molecules from bulk biomedical samples. This is especially relevant when sequencing tumors from tissue samples, as tumor sequencing is confounded by reads from non-cancerous cells including immune cells, endothelial cells, and other cells from the normal tissue [1]. This contamination is common to the majority of cancer types. For example, the reference dataset for much of cancer genomics (TCGA) is estimated to have an average purity of 62.3% across a cohort of different cancer types [2].

The ability to separate individual tumor reads from non-tumor reads in a bulk biopsy would have many useful applications. For example, in the field of early cancer detection, tumors are identified through cell-free DNA they shed [3]. More generally, the ability to extract pure tumor DNA molecules permits more sensitive detection of biological changes specific to the tumor.

In whole-genome sequenced bulk tumor biopsies, reads containing clonal somatic mutations can be confidently identified as originating from the tumor. However, the frequency of these mutations across the genome is typically too low (on the order of ∼1/Mb [4]) to be used to classify the origin of the vast majority of reads. In contrast, tumor cells typically have a more extensively altered epigenetic landscape, including changes to 5-methylcytosine at CpG dinucleotides [5]. CpG methylation patterns on individual reads therefore provide a promising basis for detection of a much higher proportion of tumor-derived reads.

A number of approaches have sought to use CpG methylation patterns to identify tumor reads both from bulk biopsies and cell-free DNA. These approaches first establish differentially methylated regions in tumors by comparing the methylation landscapes of tumors and matched normal tissue. Then, individual reads that align to marker regions of interest can be classified with a variety of algorithms. CancerDetector [6] uses a Beta-Bernoulli distribution to model the joint methylation distribution of reads within a marker region and obtain a posterior probability of a marker read having tumor origin. Other methods such as MethylBERT and DISMIR [7, 8] use machine learning approaches to classify whether reads in marker regions originate from the tumor. However, the restriction to pre-existing marker regions limits the potential sensitivity of these approaches.

New sequencing technologies from Oxford Nanopore and Pacific Biosciences can sequence substantially longer DNA molecules than Illumina’s short-read technology. PacBio HiFi typically generates reads 10-20 kb in length, while Oxford Nanopore produces reads of 10-100 kb. A single 20 kb read can capture the methylation status of an average of 190 CpG sites in the human genome, compared to just 0-5 CpG sites in 300 bp Illumina paired-end reads, although this is highly variable based on genomic location. Unlike short-read methods [9], both long-read platforms natively detect CpG methylation from DNA. Additionally, non-native marks can be detected by long-read platforms. For example, Fiber-seq uses a methyltransferase to mark open chromatin regions by methylating adenines [10]. Together, these capabilities provide a rich per-molecule epigenetic profile, making long-read sequencing a promising technology for classifying the origin of individual molecules of DNA.

In this study, we use methylation landscapes derived from deep bulk PacBio HiFi sequencing of prostate and ovarian tumors to develop ROCIT (Read Origin Classification In Tumors), a machine-learning model that can correctly identify the majority of individual tumor-derived reads from bulk biopsies across the entire genome. ROCIT does not require the use of a tissue type-matched normal biopsy, nor the explicit identification of tumor differentially methylated regions. We show that this method can enhance other sequencing capabilities by improving detection of tumor somatic mutations.

## Results

### Dataset and training

We sequenced a cohort of two prostate and four ovarian tumors as well as corresponding matched adjacent normal tissue with PacBio HiFi sequencing. This was supplemented with matched short-read sequencing for both the tumor and adjacent normal tissue with Illumina. The ASCAT copy number caller [11] determined the purity of the prostate tumors (identified as A and B respectively) as 42% and 51%. The ovarian samples A, B, C and D were determined to have purities of 90%, 46%, 83% and 77%. Notably, the short-read sequencing-based tumor type classifier CUPPA [12] classified ovarian sample B as endometrial in origin, possibly explaining its lower purity as ovarian tumors are generally expected to have high purity [13]. All other tumors were classified by CUPPA as matching the provided histological classification.

We used somatic mutation information to obtain a labeled subset of long-reads from each bulk biopsy with ground-truth tumor and non-tumor origin (Fig. 1a). All reads that contained a tumor SNV with sufficient quality (Methods) were classified as originating from the tumor. When identifying reads with a known non-tumor origin, we note that the absence of an arbitrary somatic mutation does not necessarily mean a read derives from a non-tumor cell. Therefore, we identified the set of SNVs that were present on all copies of a given haplotype in all tumor cells (Methods). As these mutations are present on all copies in a tumor, a haplotype-matched read missing such a mutation was labeled as non-tumor. We obtained an additional set of non-tumor-labeled reads by identifying regions with clonal loss of heterozygosity (LOH). Any read in such a region that phased to the haplotype that was lost in the tumor cells was classified as having non-tumor origin. In total, a mean of 1.42% (range 0.74-1.80%) of reads in each bulk biopsy could be given a label, with a mean of 35.3% (range 15.0-63.0%) of these reads identified as having tumor origin (Fig. 1b, Supplementary Table 1).

**Figure 1:**
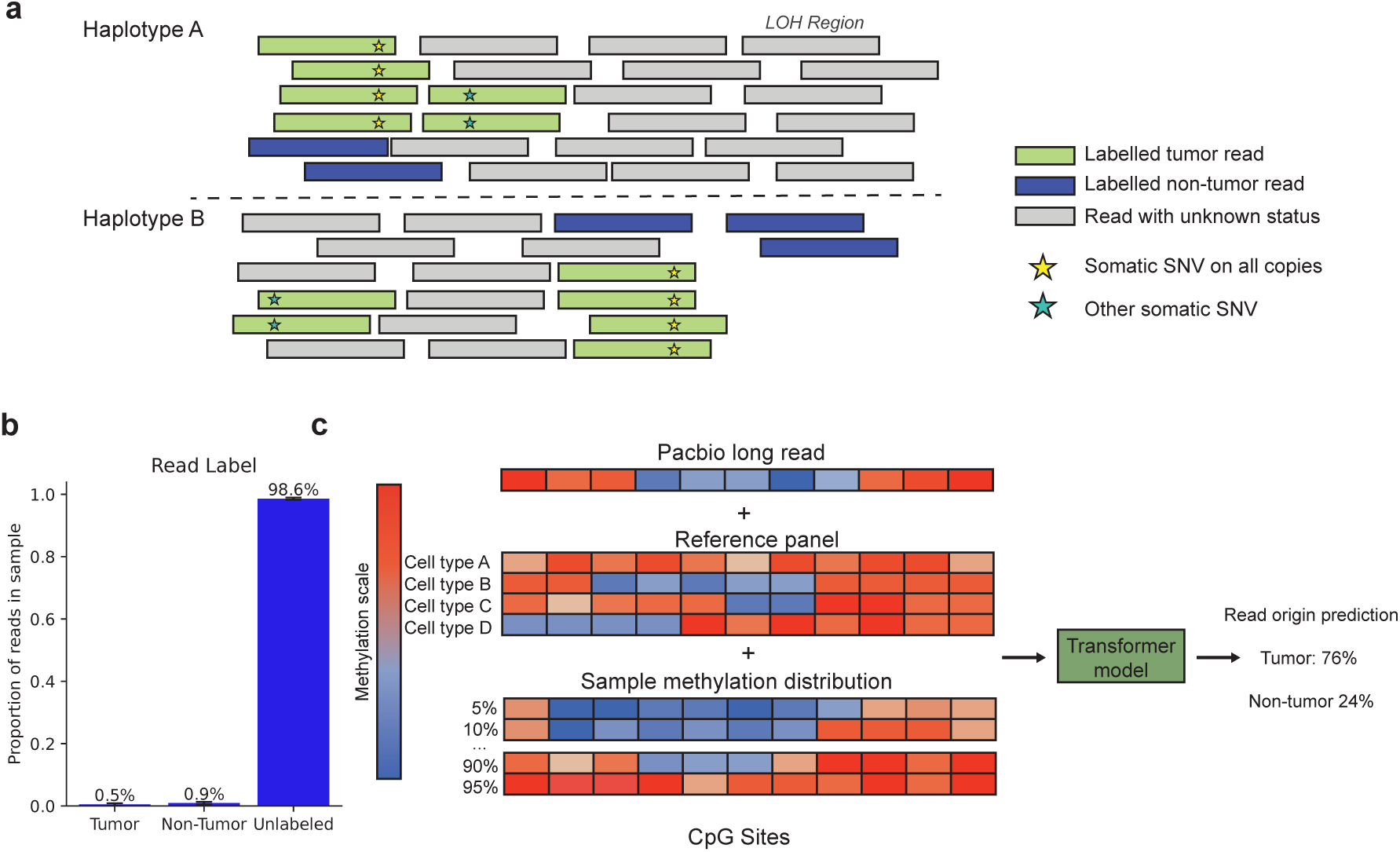
An overview of the ROCIT method and its training data. **a.** Schematic for obtaining training data reads with ground-truth tumor and non-tumor origin from somatic mutation and copy number data. All reads containing somatic mutations are labeled as tumor origin. Reads missing a clonal somatic mutation are labeled as non-tumor if they are on the same haplotype as the somatic mutation. Reads on the lost allele in regions of clonal tumor LOH (Loss of Heterozygosity) are labeled as non-tumor. **b.** Average proportion of reads per sample classified by origin as tumor, non-tumor, or unlabeled (no ground-truth label was possible). Error bars represent standard deviation across samples. **c.** Schematic of the set of features used by the model to classify each read. The methylation scale corresponds to the probability of methylation for CpG sites in individual reads for the long-read and sample methylation distribution and the average methylation for a CpG site across cells for the cell type reference panel.

This ground-truth set of labeled reads was used to train a machine learning model that classified the remaining unlabeled reads for each sample. We extracted a set of methylation-related features that the model used to classify each read. The primary feature was a vector of methylation probabilities for each CpG site in the read (Fig. 1c). We supplemented the read level methylation probabilities with other contextual features. For each CpG site contained in the read, we used cell-type-specific reference atlases from 84 different cell populations [14] (Fig. 1c). We reasoned that this would improve model performance as tumor cells have a single common ancestral cell, whereas infiltrating cells in the tumor microenvironment likely derive from a number of different cell types.

We also used the distribution of methylation probabilities for each CpG site across all reads in the bulk tumor sample as another feature (Fig 1c). This allows the model to use the relative rank of the methylation probability of each CpG site across a given read, relative to the corresponding CpG site across the rest of the reads. A binary classifier was trained to use these features to predict tumor and non-tumor origin using a small encoder-only transformer architecture [15]. We trained a separate model for each bulk tumor biopsy individually. In each tumor, reads on chromosomes 4 and 21 were held out as a validation set for hyperparameter tuning while chromosomes 5 and 22 were used for a final test-set evaluation of model performance. Using early stopping on our validation set (Methods), training finished after a median of 10 epochs, with a range of 1-11 epochs across the 6 samples in our cohort.

### Classification performance

The model demonstrated excellent discriminatory power on our test set across all samples, with a mean area under the receiver operating characteristic curve (AUC) of 0.933 (Fig. 2a, S1, Supplementary Tables 2-7). We evaluated the utility of the supplementary information used for classi-fications by training a set of models with read methylation alone, and with either the cell atlas or CpG rank information. This demonstrated that using all features together was required to obtain best performance on each of the tumor samples (Fig. 2b, S2). In particular, the model performed substantially worse when trained on individual read-level methylation values alone, demonstrating the utility of adding supplemental information about CpG sites covered by the read.

**Figure 2:**
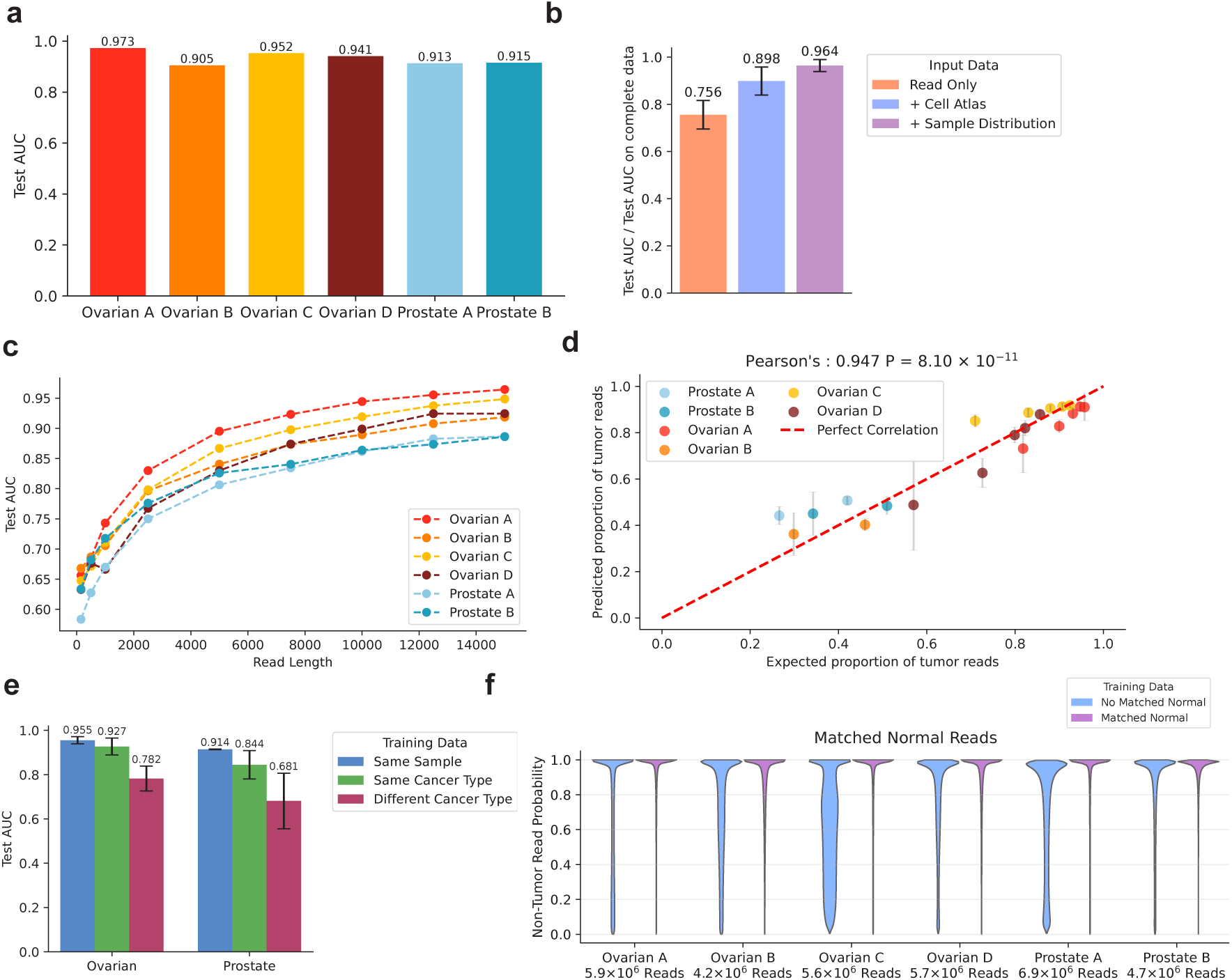
ROCIT accurately classifies reads across the genome **a.** Area under the receiver operator characteristic curve (AUC) on the test datasets. **b.** AUC results on the test datasets with different input features for classification divided by the AUC obtained from training on complete data. **c.** AUC results on the test datasets for models where input reads are cropped to a fixed length. **d.** The expected proportion of tumor reads, determined from tumor copy number and purity against the observed proportion of assigned tumor reads in the non-training set of reads from each bulk biopsy. The error bars refer to standard deviation across genomic segments with the same total copy number state. **e.** AUC results for the transformer model trained on the read data from one of the samples and applied to the test-set distribution of labeled reads from all samples. **f.** The distribution of non-tumor read probabilities for reads in matched adjacent normal bulk biopsies obtained from models where reads from the matched adjacent normal were included or excluded from the training set.

We tested an alternative classification approach by training an XGBoost [16] model on correlation coefficients of the CpG methylation probabilities of each read with the reference atlas methylation landscape of each cell type (Methods). Although we obtained good performance with XGBoost (Fig. S3), it was not as effective as the transformer-based model, with an average per-sample AUC decrease of 0.07.

We sought to test the importance of read length on model performance. We first created a subset of the labeled data with reads at least 15 kb in length. With this subset of reads, we trained a collection of models where the input data consisted of reads randomly cropped to a certain length ≤ 15 kb. We found that read length had a large effect on test-set performance, with models trained on reads cropped to 150 bp having an average test AUC of 0.64 (range 0.58-0.67), rising to an average AUC of 0.92 (range 0.89-0.96) for the same reads cropped to a length of 15 kb (Fig. 2c). This demonstrates the substantial advantage of long-read lengths for accurate tumor origin classification in arbitrary genomic regions.

To validate model performance on reads without ground-truth origin, we estimated the expected fraction of tumor-derived reads in different genomic regions using local tumor copy number and purity (Methods), a feature not explicitly provided in the training process. We observed that the proportion of reads classified as having tumor origin in each genomic region was well correlated with the proportion expected from local copy number and purity (Pearson’s 0.947, P = 8.1 x 10^-11^) (Fig. 2d).

We evaluated the ability of the models to generalize to unseen tumors by applying a model trained on a given tumor to the test-set distributions of all the other tumors. The models performed less well when applied to tumors of a different type, with an average decrease in AUC per sample of 0.22 (Fig. 2e, S4). Overall performance proved much more robust when the models were applied to the same tumor type. When ovarian sample B was excluded, as it was likely endometrial as classified by the orthogonal, non-methylation-aware CUPPA algorithm, the average reduction in test-set AUC was only 0.02 between samples of the same type. Indeed, the test-set performance of ovarian sample C slightly improved when it was classified using a model trained on ovarian sample A (Fig. S4). This suggests that the patterns the model learns are generally transferable within cancers of the same type. Changes in sequencing chemistry and methylation caller between the two cohorts (Methods) likely also impacted model transferability.

We tested the models’ ability to correctly identify reads from matched normal tissue as non-tumor (Fig. 2f). The average non-tumor origin probability for reads across the cohort of matched normal samples was 69.6% (SD = 6.9%). Supplementing the model training with a small number of reads from the matched normal sample as labeled non-tumor reads (Methods) increased the average non-tumor probability for the matched normal reads to 89.1% (SD = 2.3%) (Fig. 2f), with only a small reduction in test-set classification performance on the bulk tumor biopsy (Fig. S5).

After training, we applied each model trained on a specific tumor to predict the tumor origin probability for all reads in the bulk biopsy. Each biopsy contained an average of 1.1 x 10^7^ reads (SD = 1.1 x 10^6^). We sought to demonstrate the potential of ROCIT to enhance the capabilities of several bioinformatic applications, including somatic variant calling and identifying epigenetic changes in the tumor, independent of the epigenetics of infiltrating non-tumor cells.

We sought to identify CpG sites with the greatest influence on predicted tumor origin probability. Following earlier interpretability approaches [17, 18], we applied a gradient descent approach with a sparsity penalty (Methods) to modify the methylation probabilities for a small number of CpGs in each read to reverse its original classification. Reads that were classified as tumor were perturbed to be reclassified as non-tumor, and vice versa (Fig. 3a).

**Figure 3:**
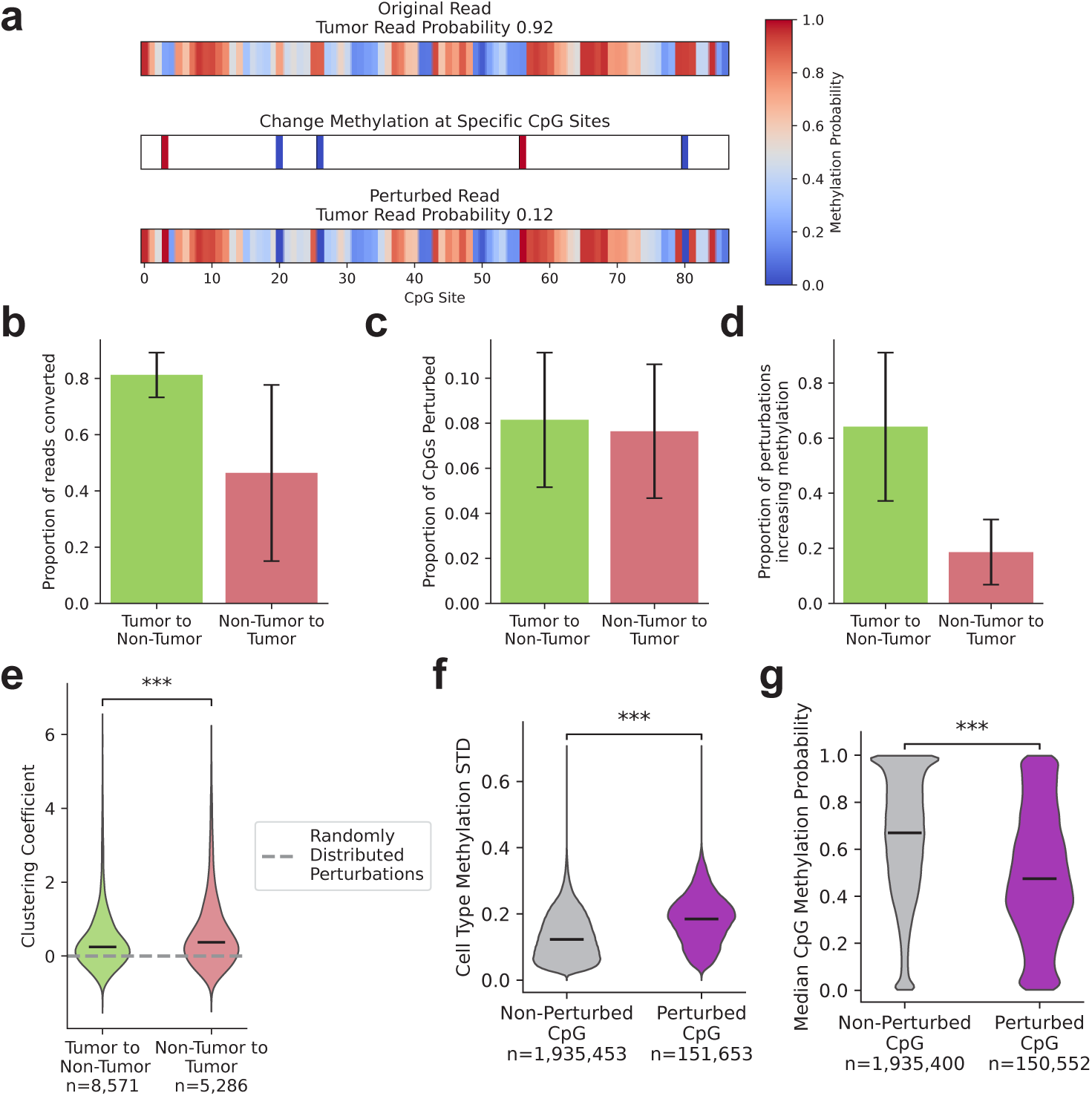
ROCIT can be interpreted by perturbing the methylation status of individual CpGs. **a.** An illustrative example of a perturbation of measured methylation probability at a limited number of sites leads to a change in ROCIT predicted tumor origin probability. **b–d.** Error bars indicate standard deviation across samples. **b.** The proportion of reads successfully converted by CpG methylation perturbations. **c.** The proportion of perturbed CpG sites in reads with successfully converted classifications. **d.** The proportion of perturbed CpG sites where the methylation probability was increased in reads with successfully converted classifications **e.** Clustering coefficients of perturbed CpG sites for reads with successfully converted classifications. Higher values indicate stronger clustering. Mann-Whitney U test, ***: *P <* 0.001 **f-g.** The standard deviation in average methylation across the cell type methylation data (**f**) and median sample-wide CpG methylation probability (**g**) for unperturbed and perturbed CpGs in successfully converted reads. Mann-Whitney U test, ***: *P <* 0.001

By applying this gradient descent approach to 23,761 reads in the test-set distribution (Methods), we were able to successfully change the classification of 58.4% of reads by perturbing an average of 8.8% of CpGs in each read (Fig. 3b-c). By varying the sparsity penalty, we could increase the proportion of successfully reclassified reads, at the cost of perturbing a higher proportion of CpGs (Fig. S6). We were able to convert a substantially greater fraction of read classifications from tumor to non-tumor (81.2%) than non-tumor to tumor (46.4%). This varies considerably by sample (Fig. S7), with the highest purity samples having the lowest rate of successful non-tumor to tumor perturbations.

We also observed a difference in the methylation directionality of the perturbations. When reclassifying reads from tumor to non-tumor, 64.1% of the perturbed CpGs had an increased methylation probability, compared with only 18.6% of CpGs in non-tumor to tumor reclassified reads (Fig. 3d). This likely reflects that tumor reads are less likely to be methylated at CpGs on average than non-tumor reads in a bulk sample. However, ovarian samples A and C had notably different behavior to the overall trend, with perturbations most frequently decreasing methylation in both reclassification directions (Fig. S8).

We observed that the perturbed CpGs were significantly closer together on the reads than expected under a model of random distribution (Fig. 3e, Methods). This was the case for reads reclassified in both directions, although the non-tumor to tumor reads had a higher average clustering coefficient.

Perturbed CpGs displayed significantly different cell reference atlas and sample methylation distributions to non-perturbed CpGs. The perturbed CpGs had a significantly higher average standard deviation of methylation probability across cell types in the reference atlas (0.185) compared to non-perturbed CpGs (0.123), as well as a lower median methylation probability in the CpG distributions for each sample (47.5% compared to 67.0%). We found that perturbations that made a read closer to CpG markers for epithelial cell types increased tumor origin probability, and perturbations that made a read closer to immune cell CpG markers increased non-tumor origin probability (Fig. S9, Methods).

### Improving somatic SNV calling

The identification of somatic tumor mutations from a bulk biopsy and matched normal tissue can be confounded by a number of factors [19]. For example, it is difficult to correctly distinguish artifacts and sequencing errors from true somatic variants, leading to imperfect variant calling. We sought to test the potential of ROCIT to improve SNV calling in long-read sequencing. We reasoned that true somatic SNVs would likely have a higher proportion of variant-containing reads labeled as tumor origin compared to artifacts and errors (Fig. 4a).

**Figure 4:**
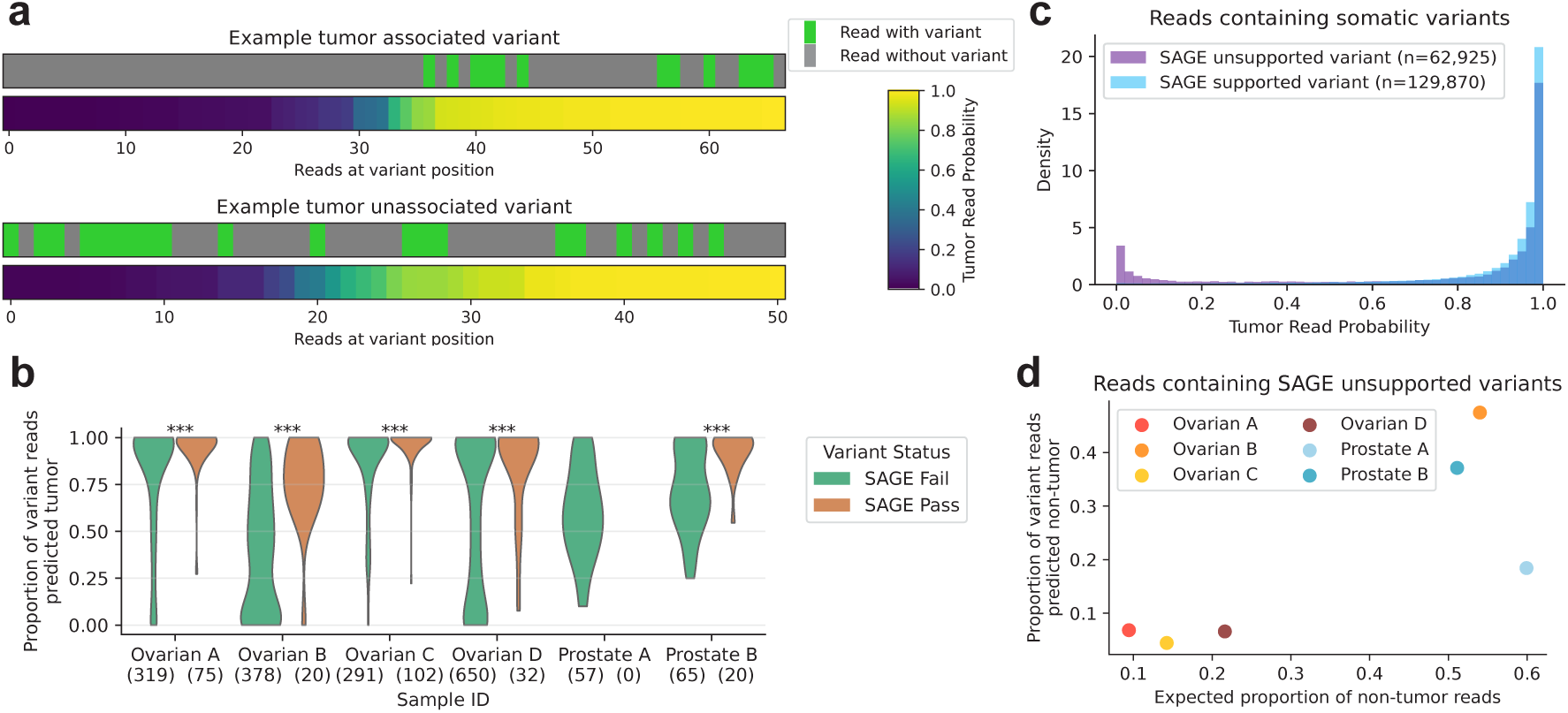
ROCIT enables more accurate SNV calling for acquired somatic tumor variants. **a.** Matched variant statuses of reads covering a given position and predicted tumor probabilities for the read. **b.** The proportion of long-reads containing variants from the set of somatic SNVs identified by the short-read SAGE caller, split by SAGE filter status. All SNV positions tested were not called as a variant in the long-read data. Mann-Whitney U test, ***: *P <* 0.001, **: *P <* 0.01. **c.** Tumor-origin probabilities for reads on non-training chromosomes containing somatic SNVs identified by DeepSomatic on the long-reads. Distributions split by whether each variant was identified as a high-quality variant by SAGE using short-read sequencing. **d.** Proportion of reads containing SNVs with DeepSomatic PASS status but without short-read SAGE support against the proportion of reads in a sample expected to come from non-tumor cells.

While many approaches are available to scan for somatic SNVs, we demonstrate ROCIT’s potential by using a set of genome positions where callers likely missed true somatic variants. These positions were found by collecting SNVs identified by the SAGE short-read caller in the matched short-read sequenced samples. We then restricted to SNVs that had supporting reads in the long-read sample but were not identified by DeepSomatic [20] as true somatic variants (Methods). To ensure adequate statistical power for proportion estimation, we restricted our analysis to SNVs with at least ten variant-containing reads in the long-read sample. As a control, we also analyzed a matched set of short-read SNVs that failed quality control checks by SAGE and were also found in ten reads but not called as somatic mutations in the long-read biopsy for each tumor.

In every sample where they were observed, SNVs that passed SAGE validation had a significantly higher proportion of tumor origin reads than those that failed (Fig. 4b). Tumor purity strongly influenced this effect, as purer samples naturally contain a higher baseline proportion of tumor-associated reads. This is evident in the lower purity samples, with ovarian B and prostate B having substantially larger differences between SAGE pass and fail SNVs in their tumor-assigned read proportions.

Our results can also be used to identify variants called by somatic variant callers that are likely not true somatic variants. The tumor origin probability distribution for reads containing DeepSomatic-called SNVs on non-training chromosomes exhibits a large tail of low predicted probability (Fig. 4c). These SNVs are disproportionately unsupported by SAGE in the short-read sequenced samples, suggesting that they may not be true somatic SNVs. Again, this effect is dominated by the lower-purity ovarian B and prostate B samples (Fig. 4d, S10).

### Read-level methylation and chromatin accessibility landscapes

Long-read sequencing enables the rich analysis of genetic and CpG methylation changes several kilobases in length at the level of a single molecule. This analysis can be further enriched with Fiber-seq [10], where methylation of adenines by the non-specific methyltransferase Eco-GII enables the detection of regions of open chromatin. ROCIT then enhances these analyses by distinguishing changes specific to tumor and non-tumor cells. We demonstrate this potential of ROCIT by considering an example, the *PTPRT* gene in ovarian sample B.

*PTPRT*, a known tumor suppressor in many cancer types [21, 22, 23], has a complex regulatory pattern with a long non-coding RNA upstream of its transcription start site. We observe substantial alternative regulation at this locus, with reads assigned to the tumor showing a strong methylation shift. Approximately 10 CpGs become methylated closer to the promoter in tumor reads, and approximately the same number of CpGs located a further 1000 bp downstream also become demethylated (Fig. 5a). This is accompanied by the opening of a FIRE element (Fiber-seq Inferred Regulatory Elements) identified by Fiber-seq in the reads predicted to originate from tumor cells specifically. Within the gene body of *PTPRT*, we observe tumor-specific demethylation combined with a FIRE peak of open chromatin starting approximately 1500 bp downstream of the *PTPRT* promoter, suggesting that the gene expression had been reduced, since gene body methylation is typically a signal of transcriptional activation [24].

**Figure 5:**
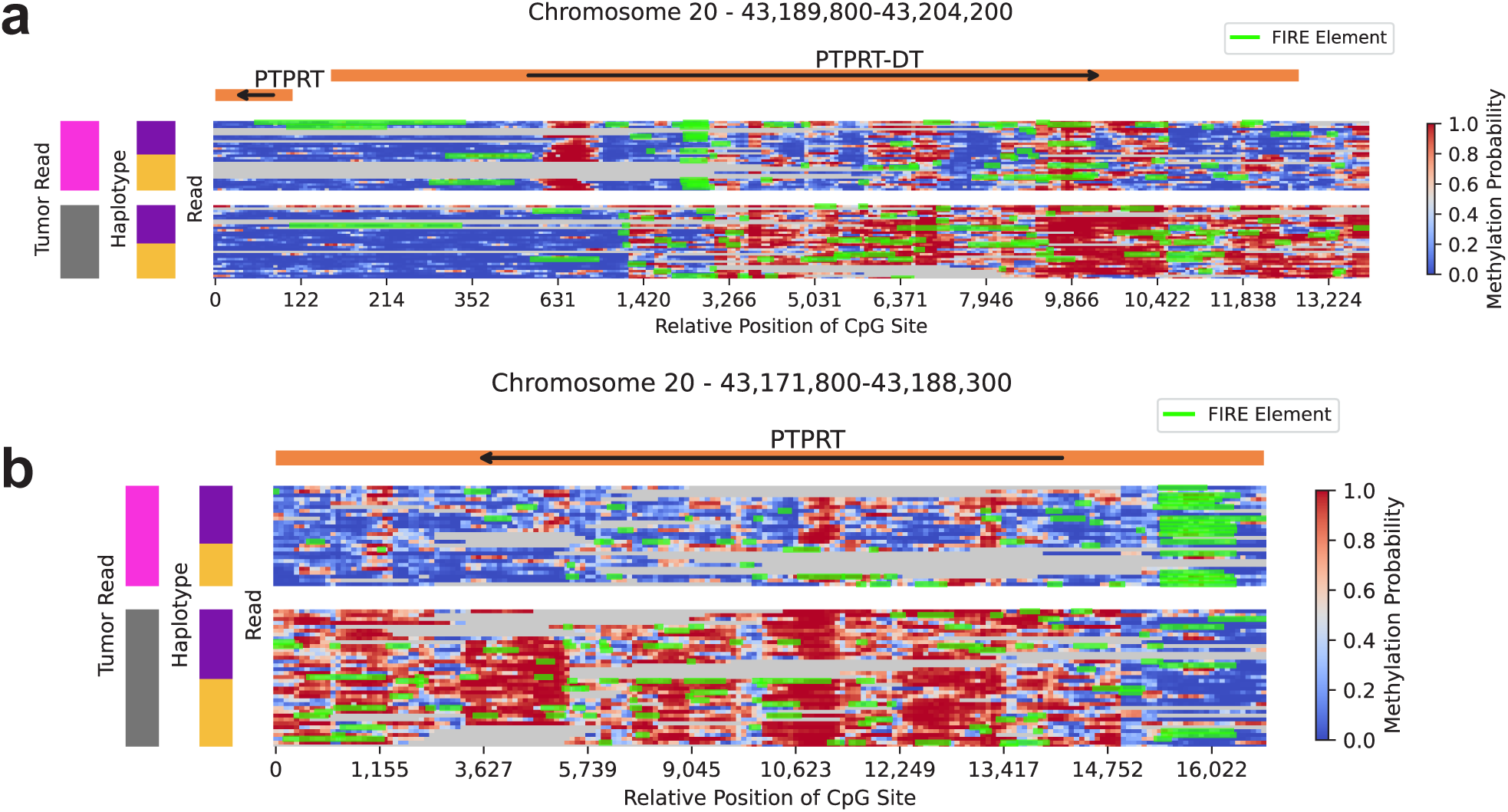
Read-level methylation and chromatin accessibility landscapes. Read-level methylation and chromatin accessibility at the PTPRT locus in ovarian sample B. **a.** *PTPRT* Divergent Transcript (*PTPRT-DT*). **b.** Section of *PTPRT* gene body. Columns represent individual CpG sites; rows represent one or more reads, as non-overlapping reads with the same tumor-read and haplotype status share rows. Gray denotes uncovered CpG positions. Green marks Fiber-seq Inferred Regulatory Elements (FIREs). Arrows indicate transcription direction. Sidebars denote tumor (pink) versus non-tumor (gray) reads and arbitrary haplotype assignments (purple, yellow).

## Discussion

Long-read sequencing platforms provide an unprecedented opportunity to measure the joint genetic and epigenetic information for individual DNA molecules 10 kb and greater in length. We show that this technology provides sufficiently rich information to be able to identify the cellular origin of individual DNA molecules. We demonstrated with ROCIT that this information can be used to accurately classify the majority of reads in a bulk sample with a tumor or non-tumor origin.

Notably, ROCIT makes no assumptions about the methylation distribution of non-tumor cells in the biopsy. This contrasts with other methods for methylation deconvolution that require matched adjacent normal tissue to use as a proxy for non-tumor cells [25]. Although all of the tumor samples in our cohort had matched adjacent normal tissue samples, other sources of matched non-tumor DNA, such as peripheral blood mononuclear cells, would work equally well, as non-tumor tissue is only required for distinguishing germline and somatic variants.

We demonstrated ROCIT on a cohort of six tumors sequenced with PacBio HiFi technology. However, given that we only require individual base calls and native DNA methylation identification, our approach should work equally well on bulk tumors sequenced with Oxford Nanopore long-read data. In fact, the longer sequencing lengths possible with the Nanopore platform (*>* 50 kb) [26] may lead to higher classification accuracy due to the greater information content available per individual read.

Currently ROCIT uses the CpG methylation distribution of each read, cell type-specific methylation patterns and sample-wide methylation distributions of each read for classification. It is likely that additional context will improve tumor/non-tumor classification, as well as cell type classification. For example, read sequence context and chromatin accessibility information from Fiber-seq could straightforwardly be integrated into this framework as additional features.

ROCIT models trained on one tumor performed well when applied to other tumors of the same cancer type, demonstrating that pre-trained ROCIT models can be successfully applied to classify reads in tumors without matched normal tissue. While validation across a broader range of cancer types remains necessary, these results suggest that ROCIT may ultimately be applicable in diagnostic contexts. The ability to detect individual tumor reads genome-wide could permit very sensitive detection of small numbers of tumor cells in biopsies with predominantly non-malignant tissue. Training on larger cohorts of tumors of the same type would likely further improve model generalization.

The potential of ROCIT predictions to improve SNV calling suggests the possibility of recursive self-improvement of the training process. An improved set of tumor read predictions may be achievable by using the predictions from an initial round of training to obtain a higher-quality set of variants. These variants could then be used to create a higher-quality set of training data to be used in a subsequent training round.

Although we have focused on the classification of reads into tumor and non-tumor origins, more advanced cell-of-origin classification for individual reads is possible. A recent method has developed a cell type classification from long-reads using a likelihood framework [27] and cell type methylation atlases [14]. The ROCIT methodology can likely be extended beyond tumor/non-tumor biology and could be used to identify specific cell types, as well as other cell states. Overall, we imagine that molecule-level deconvolution may open broad new capabilities, and we anticipate that these approaches may intersect well with single-cell methods for cell characterization.

## Acknowledgments

We would like to thank Jonas Demeulemeester and Maxime Tarabichi for useful discussions regarding dataset labeling and model training.

## Code Availability

ROCIT is available at https://github.com/tobybaker/rocit. Code for all analyses presented in this manuscript is available at https://github.com/tobybaker/rocit-manuscript-analysis.

## Author contributions

T.M.B conceptualized and developed ROCIT under the supervision of P.T.S. D.G processed tissue samples for DNA sequencing. N.M, D.A, N.G.D and K.C carried out bioinformatic processing of the original sequencing data. T.M.B carried out the majority of the analyses in the manuscript. N.M conducted analyses related to the methylation landscape of the tumor and infiltrating normal. C.A conducted analyses related to chromatin accessibility landscapes of tumor and infiltrating normal. S.W carried out analyses relating long and short-read variant processing. J.O and H.W worked on tumor differential methylation and chromatin accessibility landscapes. J.L. and B.K. supplied samples and expertise in ovarian tumor biology. S.L contributed to the validation of the training labels. T.M.B and P.T.S wrote the manuscript with contributions from S.L.

## Methods

### Sample handling and processing

We obtained the tissue of three prostate tumors and five ovarian tumors and matched adjacent normal tissue. The adjacent normal tissue was sourced from prostate for the prostate tumors, endometrial tissue for ovarian samples A and D, and ovarian tissue for ovarian samples B and C. One prostate tumor was removed from our analysis as somatic variant calling and copy number analysis found it to have very little to no tumor content. One ovarian tumor was also removed from our analysis as the tumor was sequenced to a lower depth in both the short (27.0x) and long-read (28.8x) samples, and had shorter PacBio reads with a median length of 11.8 kb.

All tumors and adjacent normal samples underwent whole-genome sequencing with both PacBio HiFi SMRT long-reads and Illumina short-reads. The same extracted DNA was used for long-read and short-read-sequencing for the ovarian samples. The extracted DNA was lost during the long-read sequencing of the prostate samples and so new DNA was extracted from the prostate tumor and adjacent normal tissue samples for short-read sequencing.

We obtained an average sequencing depth of 53.9× (SD = 4.4) for tumors and 27.2× (SD = 6.1) for matched normal samples on the PacBio long-reads. The average read length across all samples was 15.5 kb (SD = 2.2 kb). Due to new technology becoming available over the period of sample collection and sequencing, prostate tumors were sequenced using the original HiFi chemistry and the ovarian tumors with new SRPQ sequencing chemistry.

The ovarian tumor and adjacent normal samples were processed to assess chromatin accessibility using the Fiber-seq protocol [10, 28]. Briefly, nuclei were isolated from tissues and treated with EcoGII methyltransferase to label accessible DNA, after which high molecular weight DNA was extracted and sequenced according to standard PacBio protocol.

All short-read samples were sequenced with the Illumina sequencing platform. We obtained average read depths of 99.6x (SD = 10.0) and 34.4x (SD = 3.0) for the tumor and matched normal tissue respectively.

### Long-read bioinformatics preprocessing

All long-reads were trimmed and filtered according to standard HiFi processing pipelines. All samples were aligned to the GRCh38 reference genome without alternative loci using pbmm2; a PacBio-specific aligner built on top of minimap2 [29]. The prostate samples were aligned with pbmm2 version 1.14 using the HiFi preset, --min-gap-comp-id-perc 70.0 and --min-length 50. The ovarian samples were aligned with pbmm2 version 1.17 with default HiFi parameters used.

Germline variants were called for both normal and tumor samples using DeepVariant [30] version 1.9.0 using the PacBio specific model. Somatic variants were identified for each tumor sample using the DeepSomatic variant caller version 1.9.0 [20].

The germline variants were used to phase individual DNA sequencing reads using HiPhase version 1.5.0 [31]. We observed that sequencing artifacts erroneously classifed as variants negatively affected the accuracy of the phasing. Therefore, only variants from the adjacent normal tissue that were found in the COMMON dbSNP database version 151 were used for phasing. To improve sequencing coverage of both haplotypes, particularly in regions with tumor copy number loss of heterozygosity, we jointly phased the reads from the matched normal and tumor biopsies for each donor.

Tumor somatic copy number calls from long-read sequencing were obtained with ASCAT v3.2 [11]. We used a genomic bin size of 20 kb and an ASPCF penalty of 300.

### Short-read bioinformatics preprocessing

The short-read sequencing samples were all processed using Oncoanalyser version 2.0.0 [32], a Nextflow nf-core [33] pipeline that runs a suite of tools predominantly created by the Hartwig Medical Foundation to analyze short-read whole-genome sequenced tumors. The key components of this pipeline include REDUX for read filtering, followed by alignment to GRCh38 using BWA-mem. Small germline and somatic variants were identified with SAGE. Somatic copy number profiles were called using AMBER, COBALT and PURPLE.

### SNV subclonal clustering

We use DPClust version 2.2.8 [34] to perform subclonal clustering on the somatic variants. Somatic variant calls and read counts were obtained from DeepSomatic calls on the long-read sequencing data. The copy number for DPClust was obtained from ASCAT calls on the long-read data. DPClust was run for 10,000 iterations after 1,000 burn-in steps using a concentration parameter of 0.01.

A cluster was identified as the clonal cluster if it had a cancer cell fraction (CCF) of between 0.9 and 1.1 and was assigned at least 30% of the somatic variants.

For all samples, only SNVs with a PASS status for both DeepSomatic long-read variant calling and SAGE short-read variant calling were included in the set of SNVs that were clustered by DPClust.

### Obtaining training labels

Broadly, all reads containing a high-quality somatic SNV were classified as tumor reads. To obtain a set of known non-tumor reads, we identified mutations that were present on all copies of a haplotype in the tumor cells. Then, reads covering the region of these mutations but without the mutations itself were labeled as non-tumor reads. We obtained an additional set of labeled non-tumor reads by identifying reads on the lost haplotype in regions of tumor loss of heterozygosity (LOH).

Using these general principles, we applied a set of quality controls to ensure that only high quality variants and genomic regions are used for training labels.

### Obtaining mutation multiplicity and cluster assignments

For the ovarian cohort, SNV cluster assignments are obtained directly from DPClust, and the maximum likelihood estimate of the number of allelic mutations, its multiplicity, is obtained from DPInput, the preprocessing tool for DPClust.

For the prostate cohort, as not all of the mutations were used in clustering, the cluster and multiplicity for each variant were then assigned using a maximum binomial likelihood based on the positions and sizes of clusters obtained from DPClust. This was achieved using the expected variant fraction of a variant *f*, given the total tumor copy number *N_T_*, normal copy number *N_N_*, mutation multiplicity *n*, and cluster cancer cell fraction *c* [35].

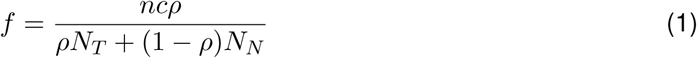

The binomial likelihood of the alternative read counts given *f* is then multiplied by the size of the corresponding cluster to obtain the overall multiplicity likelihood. The likelihood of each cluster assignment is calculated by summing the likelihoods of all possible multiplicities for a cluster.

We assumed that the range of possible multiplicity states for mutations in a clonal cluster ranged from one to the major copy number. For subclonal clusters, we assumed the mutation could only have a multiplicity of one [35].

### Variants for tumor read labels

The somatic variants then had to pass a set of general filters to be used as a tumor read identifier:

- Each SNV must be located on a canonical autosome or chromosome X.
- For the ovarian cohort, the SNV must be found in both DeepSomatic long-read and SAGE short-read somatic variant calls with PASS status.
- For the prostate cohort, the SNV must have a DeepSomatic long-read PASS status and not be filtered as a germline variant in SAGE short-read variant calls.
- The SNV must be in a HiPhase haploblock of at least 500,000 bp in size.
- There must be at least three SNV-containing reads found in the long-read-sequenced tumor sample.
- The long-reads covering the SNV position must derive from both haplotypes and all of the variant containing reads must be assigned to a single haplotype by HiPhase.
- The SNV must be assigned to a mutation cluster with a cancer cell fraction of less than 1.1.

A less stringent short-read mutation filtering step was applied to the prostate samples as the corresponding short-read sequencing was obtained from a separate tissue sample to the long-read sample.

We then applied a further filter for the phasing to ensure that that the local read phasing was consistent with the assigned copy number. Given a local tumor total copy number of *N*_Total_, tumor minor allelic copy number of *N*_Minor_, purity *ρ* and normal copy number of *N*_Minor_, the expected proportion of reads deriving from the minor allele *f*_minor_ can be calculated as:

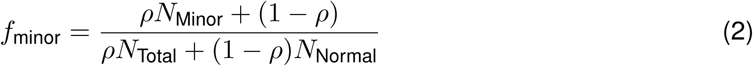

We then tested the hypothesis that the phasing counts in the phase block are distributed as ∼ Binomial(Coverage*, p* = *f*_minor_) with a two-tailed binomial test on the counts of reads assigned to the minor allele in the phase block. Reads in phase blocks with *P <* 0.1 were filtered.

Reads that contain a somatic variant and pass all of the above filters are used as the ground-truth set of reads with tumor origin.

### Variants for non-tumor read labels

To obtain an accurate set of non-tumor reads, we applied a further set of filters to the mutations that passed the tumor labeling criteria, to identify mutations that are confidently on all tumor copies of a given parental haplotype.

We identified the clonal cluster as the cluster found in DPClust for that has a cancer cell fraction between 0.9 and 1.1 and contains at least 30% of all somatic mutations for the sample. To be considered as being on all copies, a variant must have a maximum likelihood assignment to the clonal cluster.

Each variant was assigned to the major or minor allele based on whether the the haplotype that contains the variant has more reads at the position than the haplotype without the variant. Then, we remove any variants with a maximum likelihood multiplicity less than the copy number of its corresponding allele.

To ensure that the variants are in regions with high phasing accuracy, we calculated the expected proportion of reads with the variant haplotype, *f*_reads_ that contain the variant if the variant was on all copies for the haplotype in the tumor:

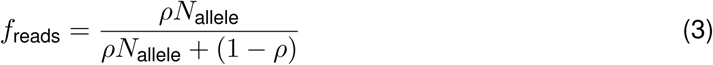

We tested the hypothesis that the proportion of reads in the variant region and with the variant haplotype that contain the variant are distributed as ∼ Binomial(Haplotype Coverage*, p* = *f*_reads_) with a two-tailed binomial test on the counts of reads assigned to the minor allele in the phase block. Reads at variant positions with *P <* 0.1 were filtered.

Variants that pass the filters in this section were identified as being on all allelic copies for a given mutation. Reads that contain the position of one of these variants, are on the same haplotype as the variant, but not the variant itself were given a label of non-tumor origin.

### Non-tumor read labels through LOH regions

In high purity tumors, the number of reads that can be confidently assigned as non-tumor through the variant method described above is relatively low, and significantly less than the number of tumor reads. Therefore, we also added a second method to obtain a set of labeled non-tumor reads using regions of tumor copy number loss of heterozygosity (LOH).

Only regions of the genome with an ASCAT copy number segment of at least a million base pairs in size were considered. The major copy number had to be between one and four and the segment had to have a coverage of at least 20 reads. Furthermore, reads assigned to both haplotypes had to be detected in the segment.

For LOH copy number segments that passed this set of filters, we used equation 2 adapted for the case where the minor allele copy number was zero. This results in a simplified equation for expected share of reads on the lost allele:

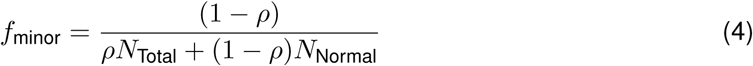

Only haplotype blocks where |*f*_observed_ − *f*_minor_| ≤ 0.05 were considered. This test was repeated on sub-intervals of each haplotype block, obtained by splitting the block into evenly sized intervals as close to 100 kb as possible. Any reads that had a maximum overlap with a sub-interval that failed this test were filtered.

All reads in haplotype blocks and genomic regions that passed these set of criteria were labeled with a non-tumor origin.

### Extracting feature sets

Four different features were given to the model to make a classification. The first two features are the measured probabilities of CpG methylation on the read and the relative reference position of each CpG site in the read. The other two features were supplemental information for the CpG sites covered by the read. We include average methylation values across cell type reference maps, as well as the distribution of CpG methylation values for the sample in question.

Only the primary alignments of reads were considered and the methylation probabilities of any CpG sites in the read without a corresponding genomic reference position were discarded.

#### Read methylation values

The probability of methylation for each CpG site in a PacBio read is provided by the Jasmine modification caller, using nucleotide kinetic data for classification. Only symmetric CpG methylation across both strands is considered as a possible state. The probabilities are provided by Jasmine as integers between 0 and 255 with a value of *M* corresponding to a probability of methylation between 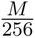 and 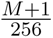. We scale these values by replacing each value with the midpoint of the corresponding bin and stacking consecutive values to form a vector of CpG methylation probabilities.

#### Read CpG positions

The relative spacing between each CpG site on the read is included as a feature using the reference position of each CpG site. The vector of sorted CpG read reference positions **R**, is scaled by the following formula:

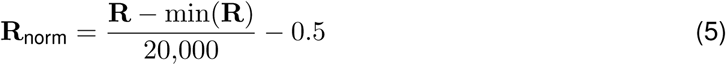

For a given read, this scaling procedure means that the values of **R**_norm_ range from -0.5 to approximately 0.5. By using a fixed denominator, we ensure that the length of the read is implicitly encoded in the relative positions, with longer reads having larger maximum values of **R**_norm_.

#### Sample methylation distribution

We represent the distribution of methylation probability across all reads for each CpG with a fixed set of percentiles. All methylation probabilities across all matching reads for a given CpG in a sample are sorted and then the 5%, 10%, …, 90%, 95% percentiles of the distribution are calculated and stored. Then for each read, the percentile values of the relevant CpGs are retrieved and an *N* × 19 matrix, where *N* is the number of CpGs in the read, is used as a feature in the model.

#### Cell type methylation distribution

For each CpG site in a given sample we obtain the cell methylation probability distribution from the whole-genome bisulfite sequenced cell reference maps provided by Loyfer et al. [14]. We downloaded the average methylation values across 28,217,448 CpGs for 84 different cell types and biological replicates from GEO dataset GSE186458 and averaged the methylation values for each CpG across the biological replicates. Then for each read, the average methylation values of the relevant CpGs are retrieved and stored in an *N* × 84 matrix, where *N* is the number of CpGs in the read.

### Model architecture

The classification model is based on the bidirectional transformer architecture [15] with an embedding dimension of 384, 6 attention heads (each with a dimension of 64) and 3 sequential transformer blocks. The transformer blocks were implemented with the TransfomerEncoderLayer from PyTorch using a feedforward dimension of 1536, layer normalization applied before attention, GELU activation [36] and no bias terms in the linear or layer normalization layers.

The input data are projected into the embedding space using a pair of dense feed-forward networks, each with two hidden layers. The first layer transforms the data from its input dimension to a dimension of 384, while subsequent layers maintain the 384-dimensional embedding space. GELU activation is used between each of the dense layers.

The first network is applied to a concatenated vector of size 21 for each CpG site. The vector is composed of the CpG methylation probability, normalized CpG reference position and methylation sample percentile distribution for the CpG. The second network is applied to the 84 cell type methylation values for each CpG site. Before projection, missing values are replaced with a learned embedding, allowing the model to infer that these entries are absent.

The two projected vectors are summed together elementwise along with a learned positional embedding for the relative position of the CpG site in the read. These set of CpG projections are prepended with a learned classification embedding vector before being passed through the transformer layers. The classification vector for each read is then projected to a classification logit with a final linear layer.

All parameters were initialized using PyTorch default values. The outputs of the absolute position embedding layer were multiplied by 0.05 to match the magnitude of the outputs of the feedforward networks.

#### Model training

A model for each sample was trained separately. For each sample, chromosomes 4 and 21 were held out for hyperparameter tuning and chromosomes 5 and 22 for a final evaluation. The maximum number of CpGs permitted per read was 511. Padding was added for reads with fewer than 511 CpGs. For the 1.9% of reads in our data with more than 511 CpGs, 511 CpGs were randomly sampled from the read at each training iteration.

The models were trained using the AdamW optimizer [37] with a learning rate of 1 × 10*^−^*^4^. A linear learning rate warm up of 100 steps was used. For regularization, the linear projection layers and transformer blocks had a dropout of 0.1 applied during training [38]. Additionally, a small amount of noise *ε* ∼ N (0, 0.01^2^) was added during training to the read methylation, sample methylation distribution and cell type methylation distribution values.

We used a weighted binary cross entropy loss function, using tumor reads as the positive class. The following positive weighting scheme was applied for balanced training:

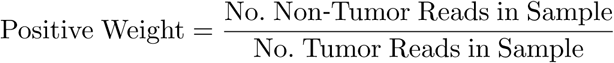

The model was trained until the area under the receiver operator curve for the tuning data had not improved for 5 epochs. At this point, training was halted and the model weights corresponding to the epoch with best validation-set performance were restored. The models were then run on the full set of reads for each biopsy. A read was classified as a tumor read if it had a tumor origin probability *p* ≥ 0.5.

#### Training with normal data

The models trained with additional reads from the adjacent normal underwent the same training process. We randomly sampled 5% of the reads from each the adjacent normal samples and gave them a label of non-tumor. These reads were separated into training, testing and validation datasets based on the same chromosome location-based criteria used for the training data derived from the tumor samples.

### XGBoost model

#### XGBoost features

The XGBoost [16] model was trained using the same set of labeled reads as was used in the transformer. We extracted a number of fixed features in order to encode the data in a format suitable for XGBoost training. To encode the overall methylation of the read, the 5%, 10%, …, 95% percentiles of the methylation probability distributions for all CpGs in the read were used as features.

The Pearson correlation coefficient between the CpG read methylation probabilities and the average cell type reference map methylation for each of the 84 cells types were used as an additional feature. We also encoded the relative difference between the methylation probability of each CpG in the read and its distribution of methylation values across all reads in the sample. We first measured the relative rank of the methylation for each CpG site against the the 5%, 10%, …, 95% percentiles of the methylation distribution of the same site across all reads in the sample. Then the 5%, 10%, …, 95% percentiles of these distribution of ranks across all CpGs in the read are used as features.

#### XGBoost training

The models were trained using the XGBoost GPU python package [16] using the histogram approximation method. As with the transformer approach, a separate model was trained for each individual sample. We applied a parameter sweep over the following set of parameters, selecting the combination for each model with the highest validation-set AUC.

**Table 1:** XGBoost hyperparameters.

| Parameter | Values |
| --- | --- |
| Max depth | {5, 7, 9, 11} |
| Learning rate | 0.01 |
| Number of boosting rounds | 15,000 |
| Subsample ratio | {0.25, 0.5, 0.6, 0.8, 1.0} |
| Min child weight | {3, 5, 10, 25} |
| Column subsample ratio | {0.25, 0.5, 0.75, 1.0} |
| Early stopping fraction | 0.1 |
| Objective | Binary logistic |
| Evaluation metric | AUC |

#### Expected share of tumor reads

For a copy number segment with purity *ρ*, total tumor copy number *N*_Total_ and normal copy number *N*_normal_, the expected share of tumor reads *f*_tumor_ for the segment is given by:

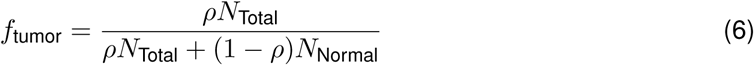

### Model interpretability

We used a perturbation approach to identify a small group of CpGs that have a large impact on tumor origin classification with ROCIT. Taking a trained model with fixed parameters, we optimized a perturbation ***δ*** applied to each read methylation vector **x** such that the predictions produced by the model *f* (**x**) are inverted, meaning that tumor predicted reads become non-tumor predicted reads and vice versa. A smooth approximation to an *L*_0_ penalty was applied to the *δ* to minimize the number of CpG sites with changed perturbations per read. Our loss function has three components:

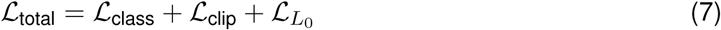

The class loss ℒ_class_ is a binary cross entropy loss between the output of the model on the perturbed methylation *f* (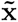) and the target binary label *y*, the inverse of the maximum probability prediction obtained from the unperturbed read *f* (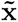).

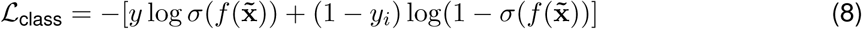

The clipping loss ℒ*_clip_*is a penalty on the perturbed methylation 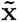 = **x** + ***δ*** to ensure that it remains within the 0-1 range.

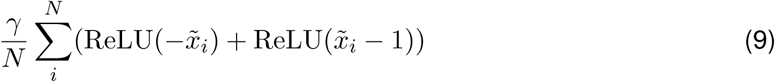

We averaged over the number of CpGs attended to in each read *N* and apply a fixed strength parameter *γ* of 50.

The ℒ_*L*_0__ loss is composed of a smooth approximation to a true *L*_0_ loss:

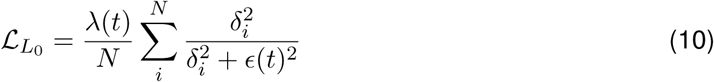

The smoothness parameter *ɛ*(*t*) and the overall strength parameter *λ*(*t*) were varied with the optimization step *t* such that the relative steepness and magnitude of the ℒ_*L*_0__ is higher at the end of the optimization process.

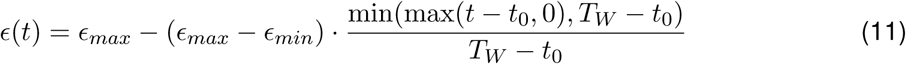

The equation for the strength parameter is similar:

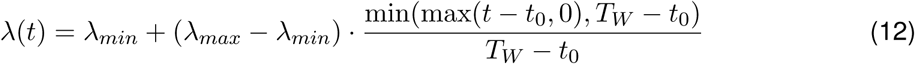

These equations are structured that the *L*_0_ penalty is only applied after the first *t*_0_ steps and then linearly increased over *T_W_*steps (Table 2).

**Table 2:**
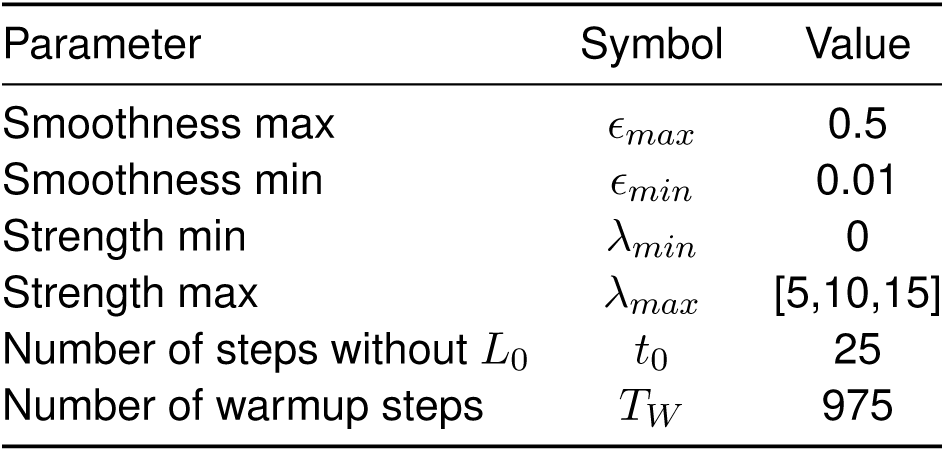
***L*_0_ penalty scheduling hyperparameters for *ɛ*(*t*) and *λ*(*t*).**

The final perturbed methylation 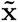 was obtained by applying an element-wise threshold to ***δ***, setting all |*δ_i_*| *<* 0.1 to 0 and then clipping 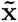 to the interval [0, 1].

We varied the strength parameter *λ_max_* over 3 different values over different optimization runs using the same reads. The perturbation ***δ*** for each read was initialized to zero. Each batch consisted of 1024 reads from a single sample. The Adam optimizer was used on the average loss across reads in the batch with a learning rate of 0.01 for *T_W_* + *t*_0_ = 1000 steps.

A total of 5,120 reads from the test-set distribution were perturbed for each sample. However, we restricted downstream analysis to reads with a predicted unperturbed probability of tumor origin *p* ≥ 80% or ≤ 20%. This left a total of 23,761 reads across the six samples in the cohort. A read was considered successfully perturbed if the predicted probability of tumor origin post-perturbation *p^′^* ≥ 80% and original probability *p* ≤ 20% or *p^′^* ≤ 20% and *p* ≥ 80%. Any CpG site with |*δ*| *>* 0.1 was considered perturbed.

#### Perturbed CpG clustering coefficients

Following the nearest-neighbor distance approach of Clark and Evans [39], we assessed whether perturbed CpG sites on individual reads were spatially clustered. For each read with a successfully swapped classification, we computed the mean nearest-neighbor distance between perturbed CpGs in CpG-index coordinates. To generate a null expectation, we permuted the identity of perturbed CpGs across all CpG sites on the read, preserving their total number, and recomputed the mean nearest-neighbor distance for each of *N* = 5,000 permutations. A clustering coefficient *C* was then defined as:

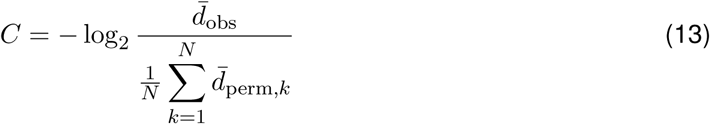

where 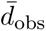 is the observed mean nearest-neighbor distance and 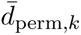 is the mean nearest-neighbor distance for the *k*-th permutation. Positive values of *C* indicate that perturbed CpGs are more clustered than expected under a null model.

#### Cell-type concordance of perturbations

We sought to test the association between perturbations at individual CpG sites and marker CpG sites for the cell types in our reference atlas. A CpG site was considered a marker site for a given cell type if the difference between the average methylation for that cell type and the average methylation across all cell types was at least 0.5. Only cell types with at least 20 marker CpGs across all successfully perturbed reads for a sample were considered.

For CpG perturbations at cell type markers that increased tumor origin probability, a cell type was considered tumor-like if the perturbation shifted methylation toward that cell type’s methylation value, and non-tumor-like if it shifted away. Conversely, for perturbations at cell type markers that decreased tumor origin probability, a cell type was considered tumor-like if the perturbation shifted methylation away from the cell type’s methylation value, and non-tumor-like if it shifted toward it.

### ROCIT somatic mutation identification cohort

We identified all SAGE somatic SNVs, independent of filter status, that had at least ten SNV reads in the long-read tumor biopsy and no reads containing the SNV in the adjacent normal long-read sequenced biopsy. We required ten reads that contained the SNV in the tumor biopsy in order to obtain sufficient tumor read prediction resolution. To eliminate any potential bias from the use of DeepSomatic variants for training data, we did not consider any SAGE variant that was within 100 bp of a DeepSomatic PacBio variant with PASS status. We calculated the proportion of reads containing each variant that were identified as tumor, using the SAGE PASS and filtered variants as our test and control cohorts respectively.

### Fiber-seq processing and analysis

6mA sites were predicted with Jasmine (v2.3.0) via the PacBio Revio Machine. fibertools [40] (v0.6.4) was then used to call nucleosome positions on the aligned sequence data. The Fiber-seq Inferred Regulatory Elements (FIRE) (v0.1.3) pipeline [41] was used to predict read-level regulatory elements and sample-level chromatin peaks. FIRE peaks were filtered to include those that pass default coverage thresholds and an FDR associated with the FIRE score less than 5%.

## Supplementary Figures

**Figure S1:**
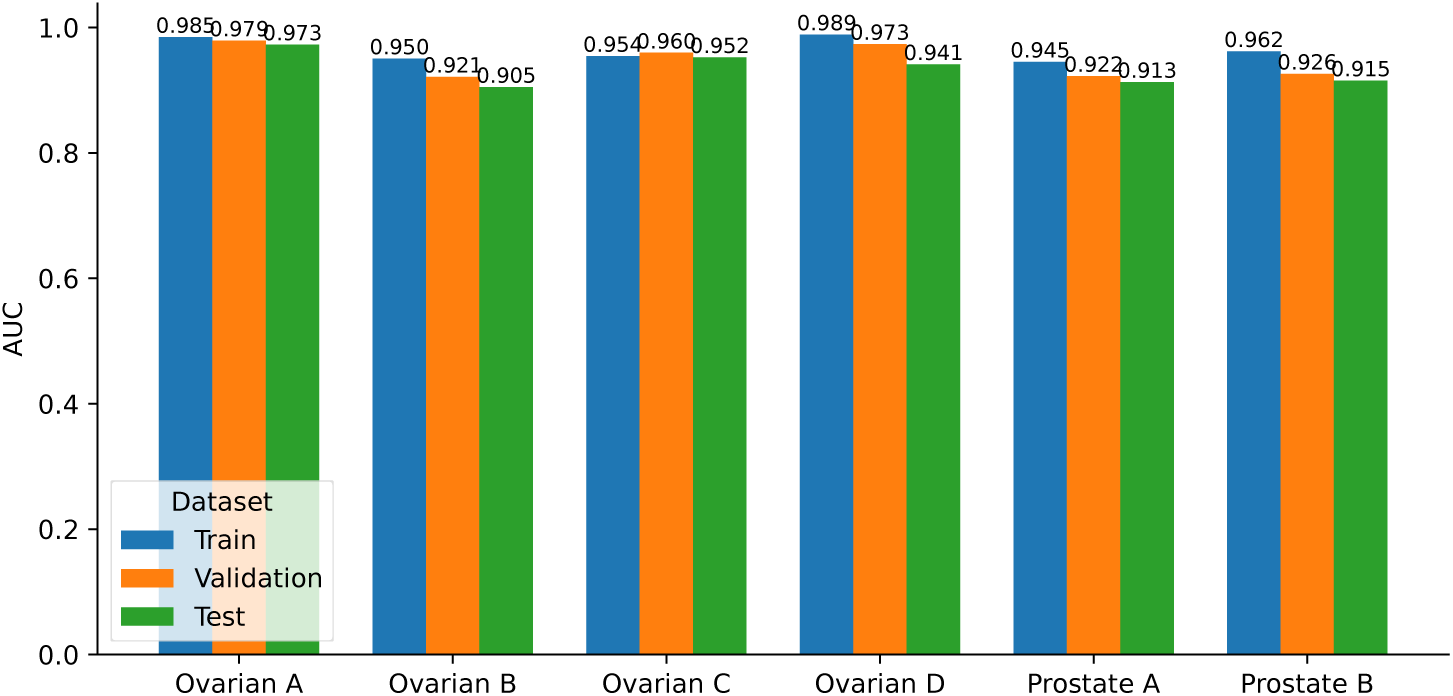
Area under the receiver operator curve results for the train, validation and test set datasets for the transformer model on the four ovarian and two prostate samples.

**Figure S2:**
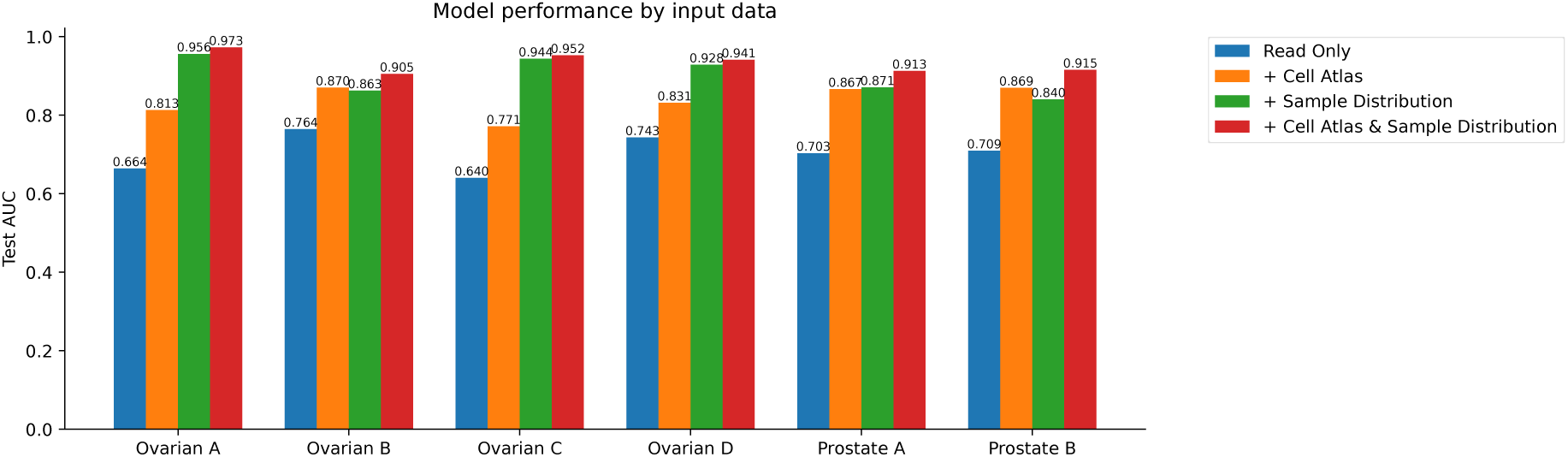
AUC results on the test datasets with different input features for classification. Results displayed for training on read level methylation alone, incorporating the CpG sample distributions and cell type methylation reference atlases separately and together.

**Figure S3:**
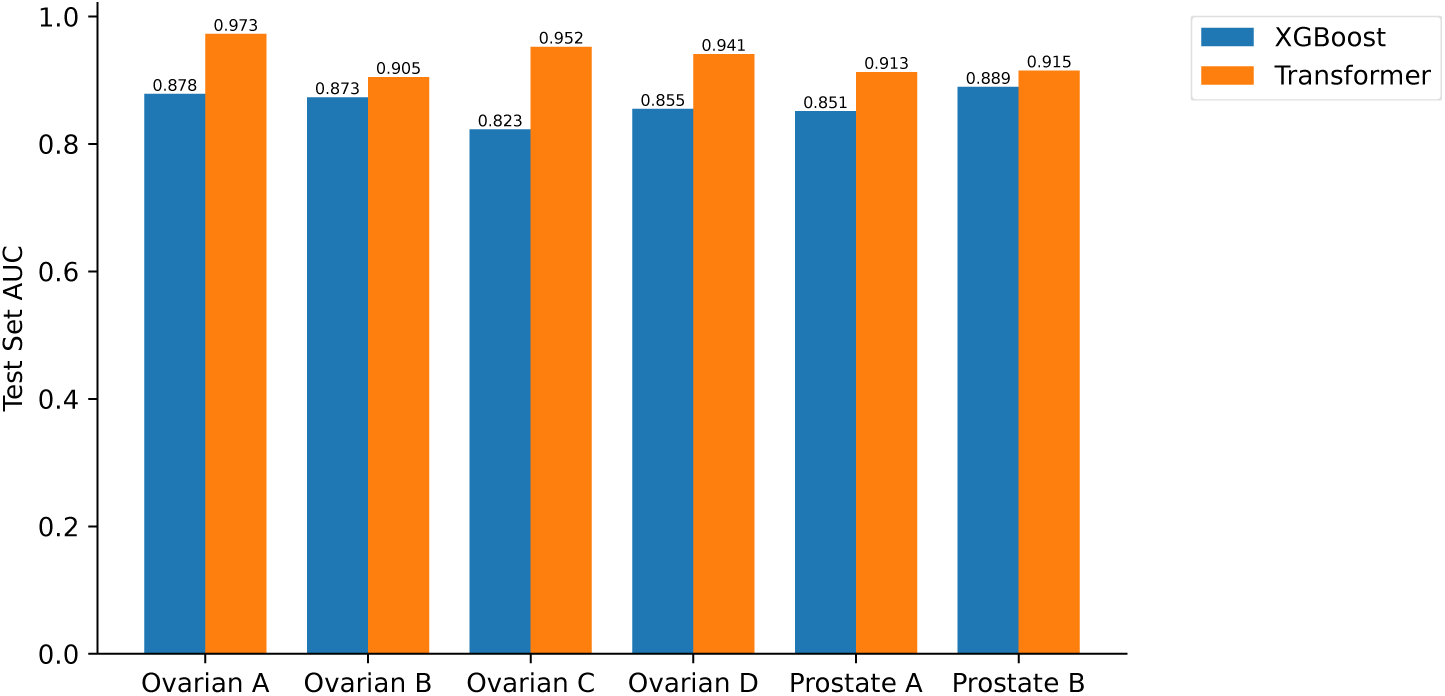
Area under the receiver operator curve results on test set datasets for the transformer and XGBoost models trained and evaluated on the same sample for the four ovarian and two prostate samples.

**Figure S4:**
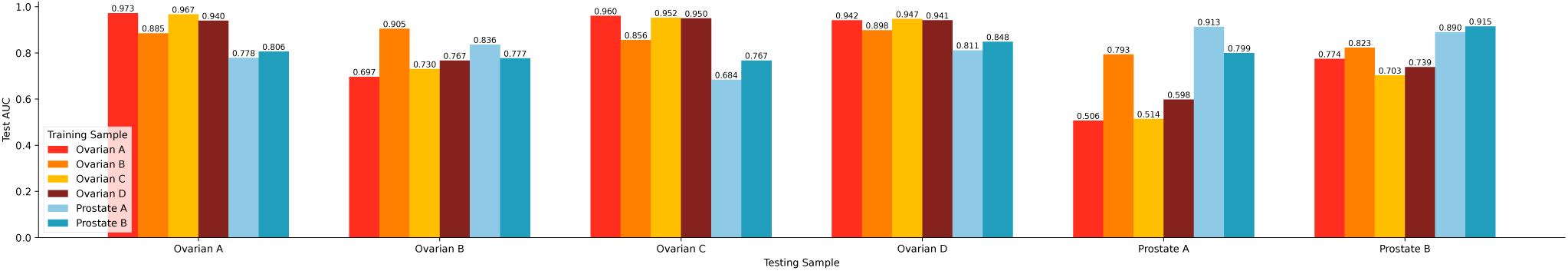
Area under the receiver operator curve results for the transformer model trained on the read data from one of the samples and applied to test set distribution of labeled reads from all samples separately.

**Figure S5:**
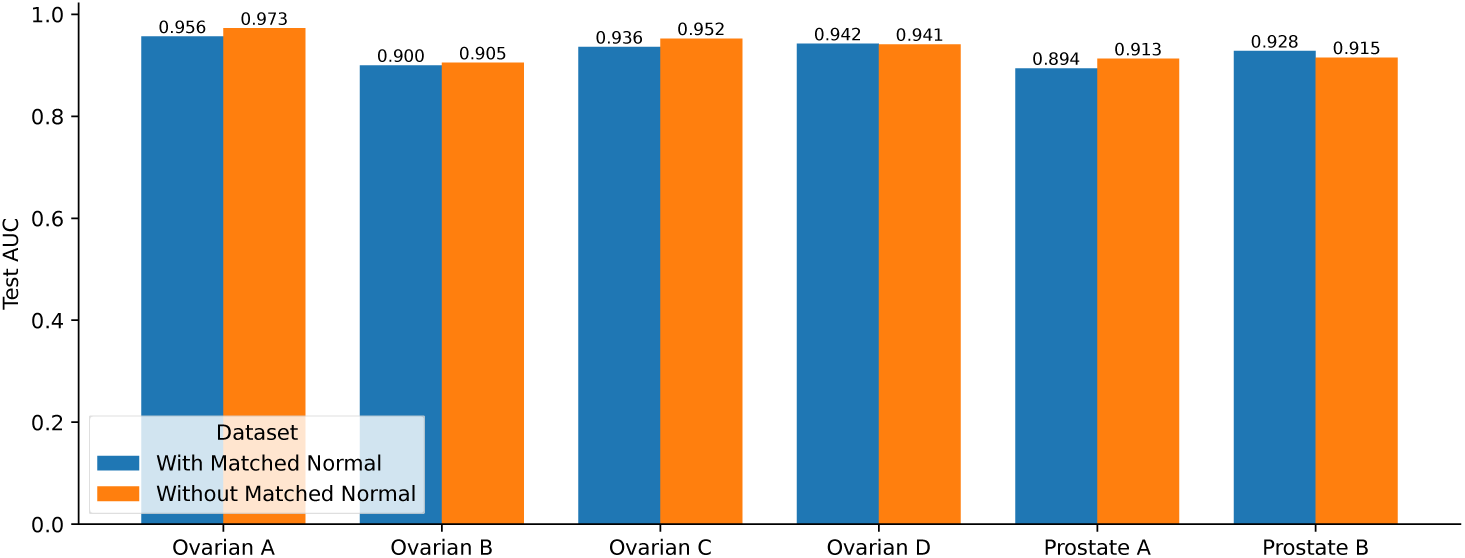
Area under the receiver operator curve results on test set datasets for transformer models with and without matched normal reads added to the training set. No matched normal reads were added to the test set.

**Figure S6:**
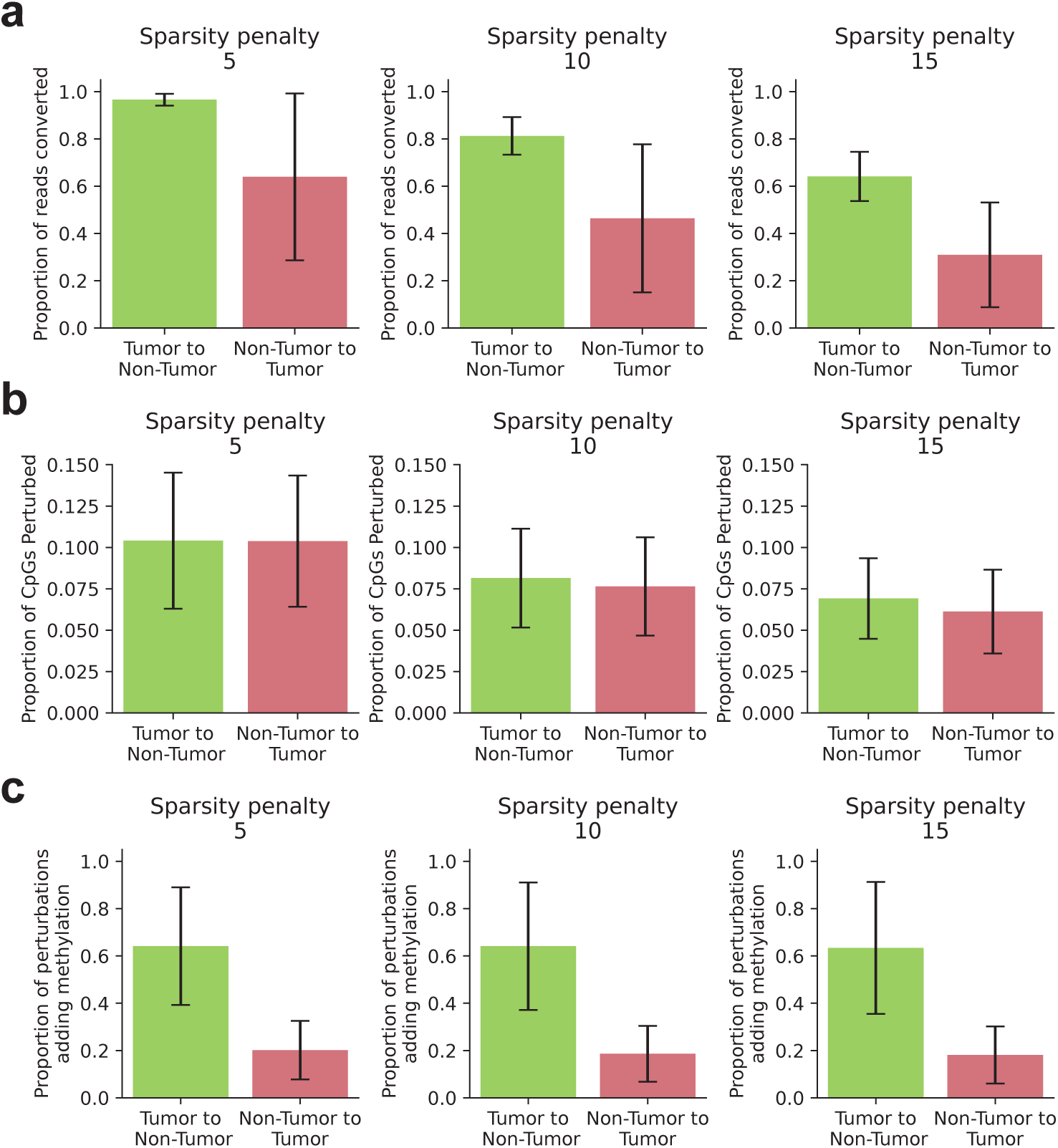
**a–c.** Error bars indicate standard deviation across samples. **a.** The proportion of reads successfully converted by CpG methylation perturbations for different *L*_0_ sparsity penalty values. **b.** The proportion of perturbed CpG sites in reads with successfully converted classifications for different *L*_0_ sparsity penalty values. **c.** The proportion of perturbed CpG sites where the methylation probability was increased in reads with successfully converted classifications for different *L*_0_ sparsity penalty values.

**Figure S7:**
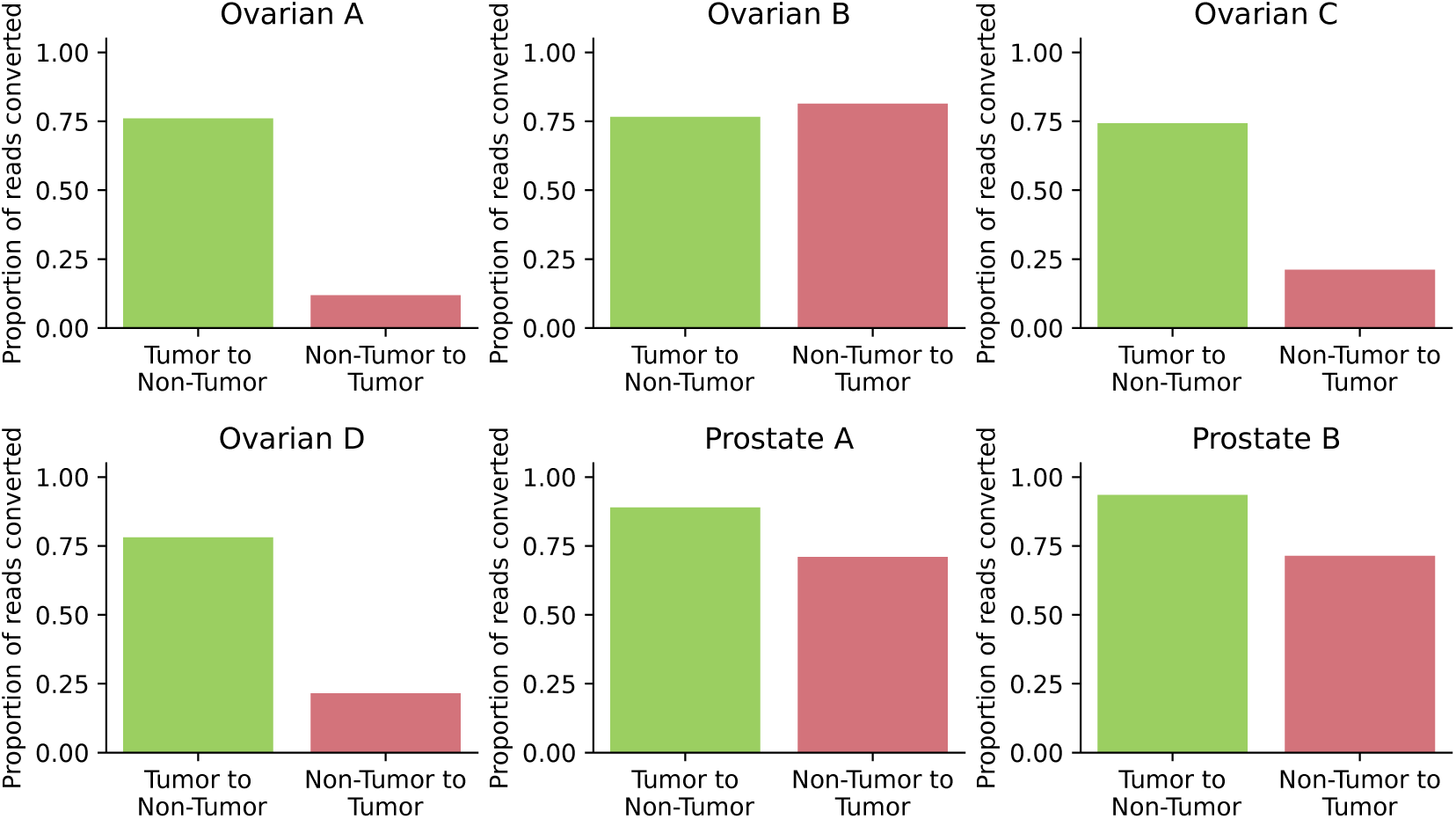
The proportion of reads successfully converted by CpG methylation perturbations split by the samples in the cohort.

**Figure S8:**
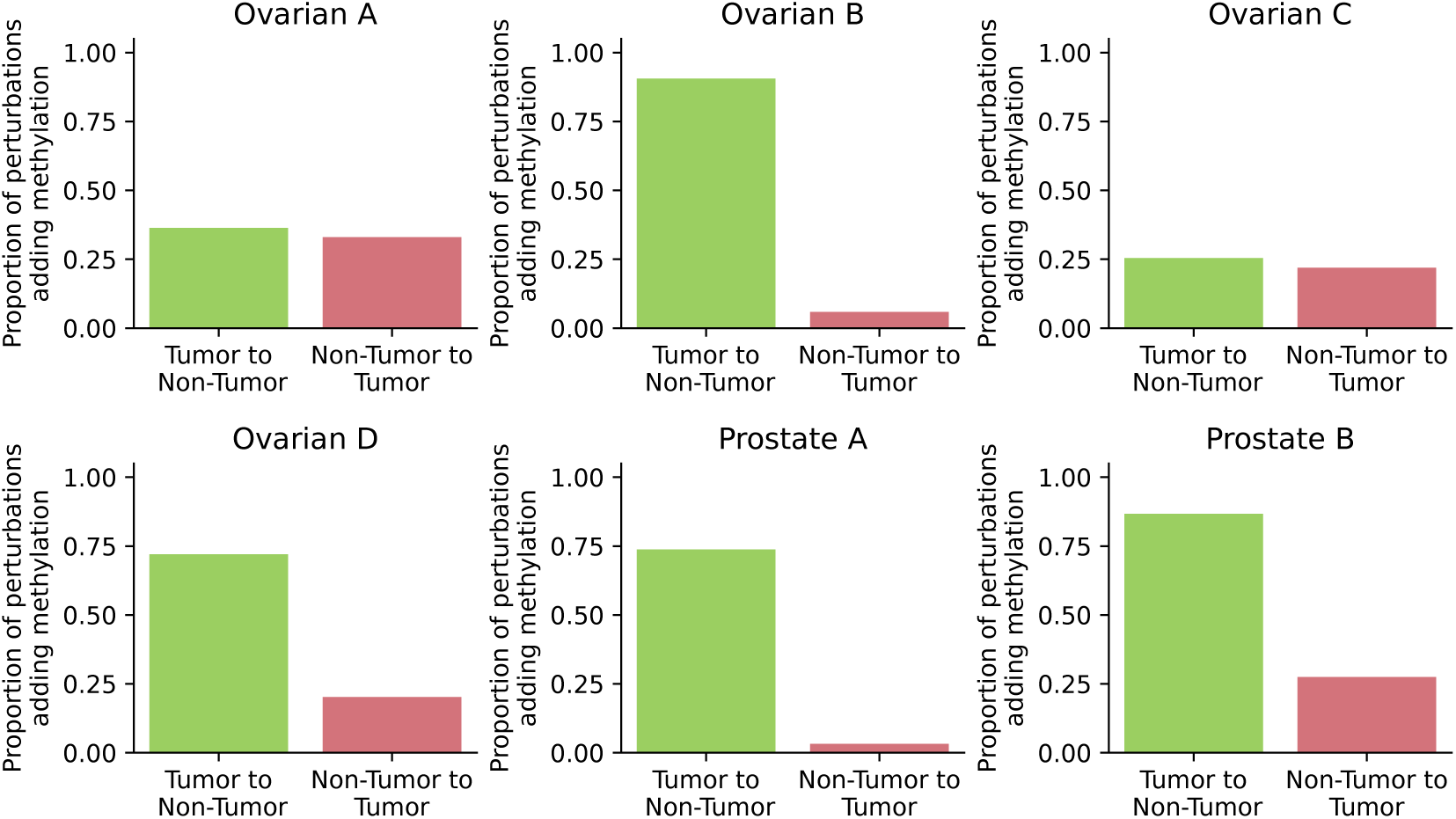
The proportion of perturbed CpG sites where the methylation probability was increased in reads with successfully converted classifications split by the samples in the cohort.

**Figure S9:**
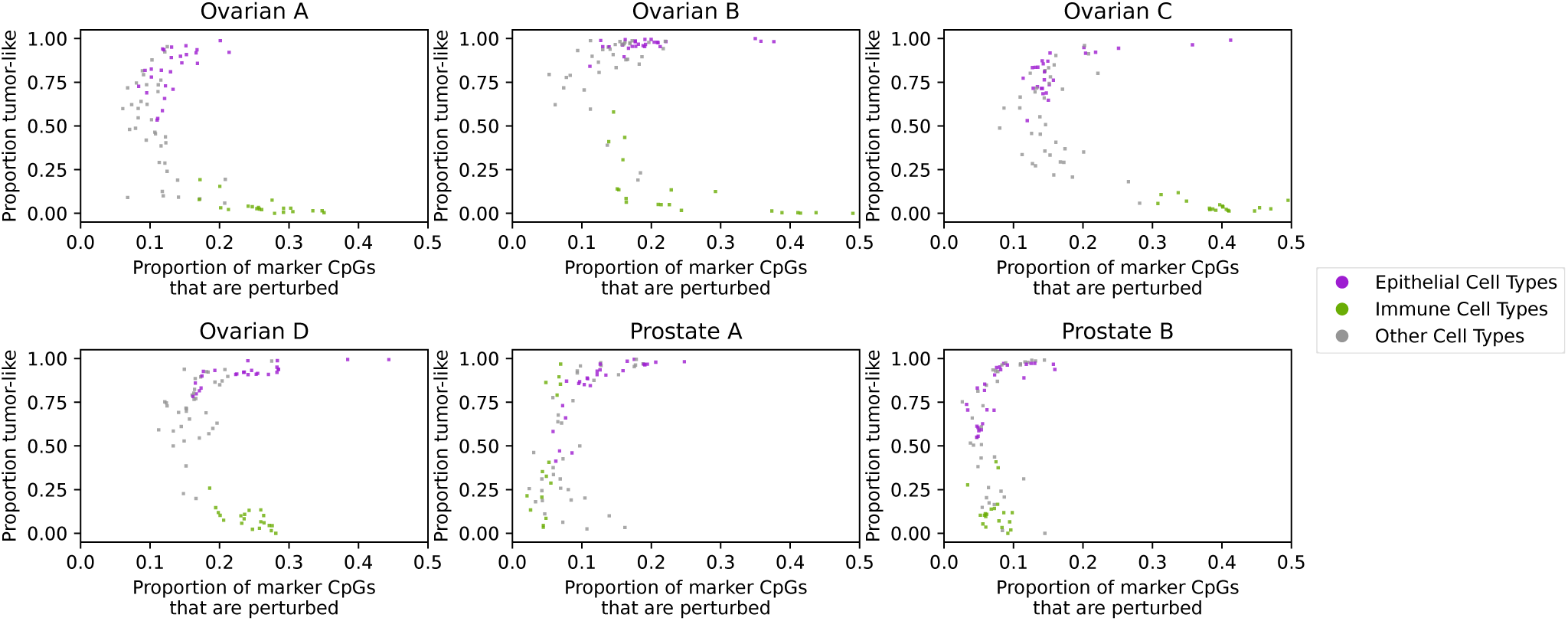
The proportion of marker CpGs for a given cell type that were perturbed in reads with successful perturbations against the proportion of perturbations for each marker CpG where the marker was associated with increased tumor origin probability for a read.

**Figure S10:**
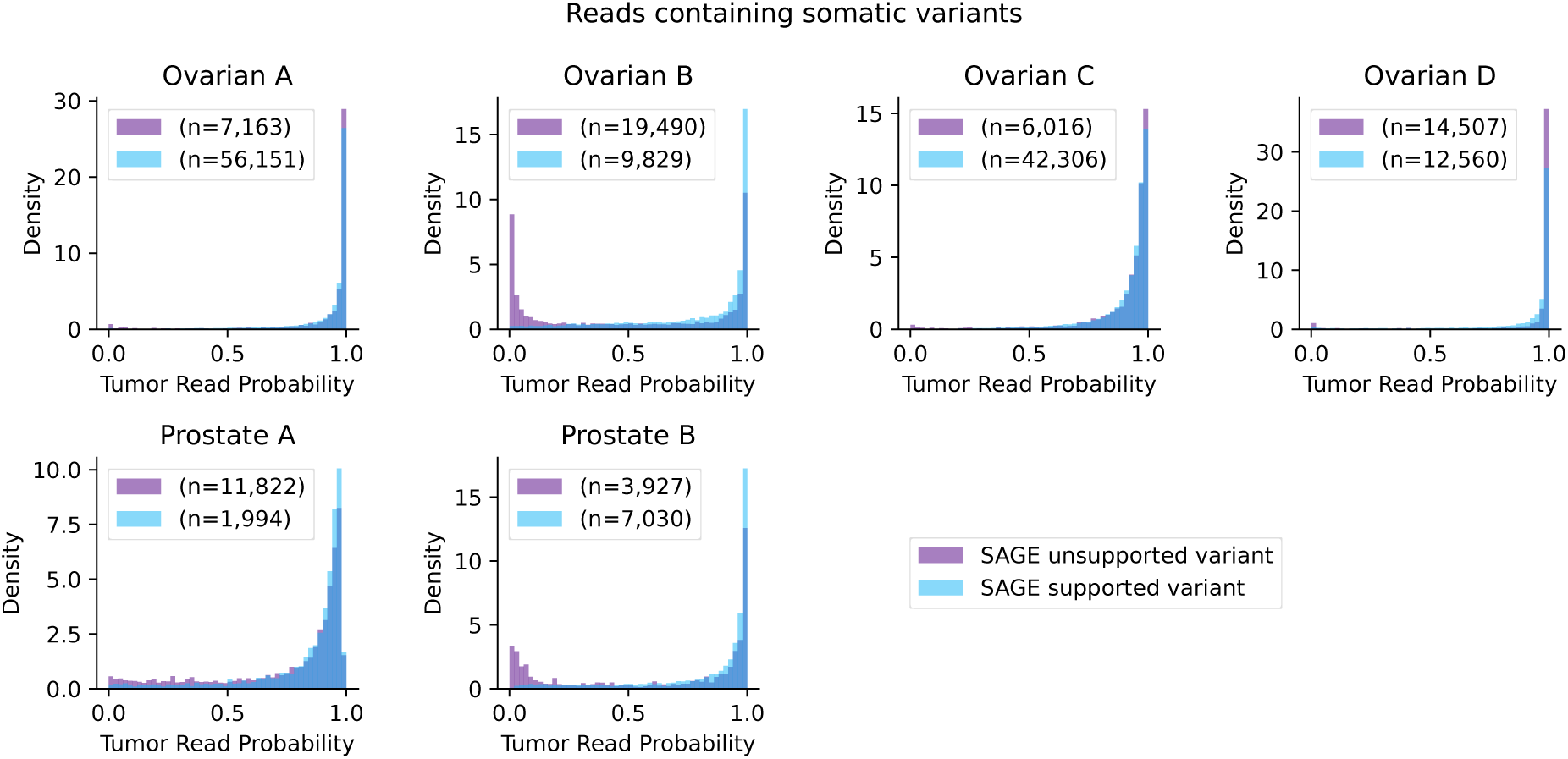
Tumor read probabilities for each sample for reads containing somatic variants identified by DeepSomatic from long read sequencing. Distributions split by whether each variant was identified as a high quality variant by SAGE using short read sequencing.

## Supplementary Tables

**Supplementary Table 1:**
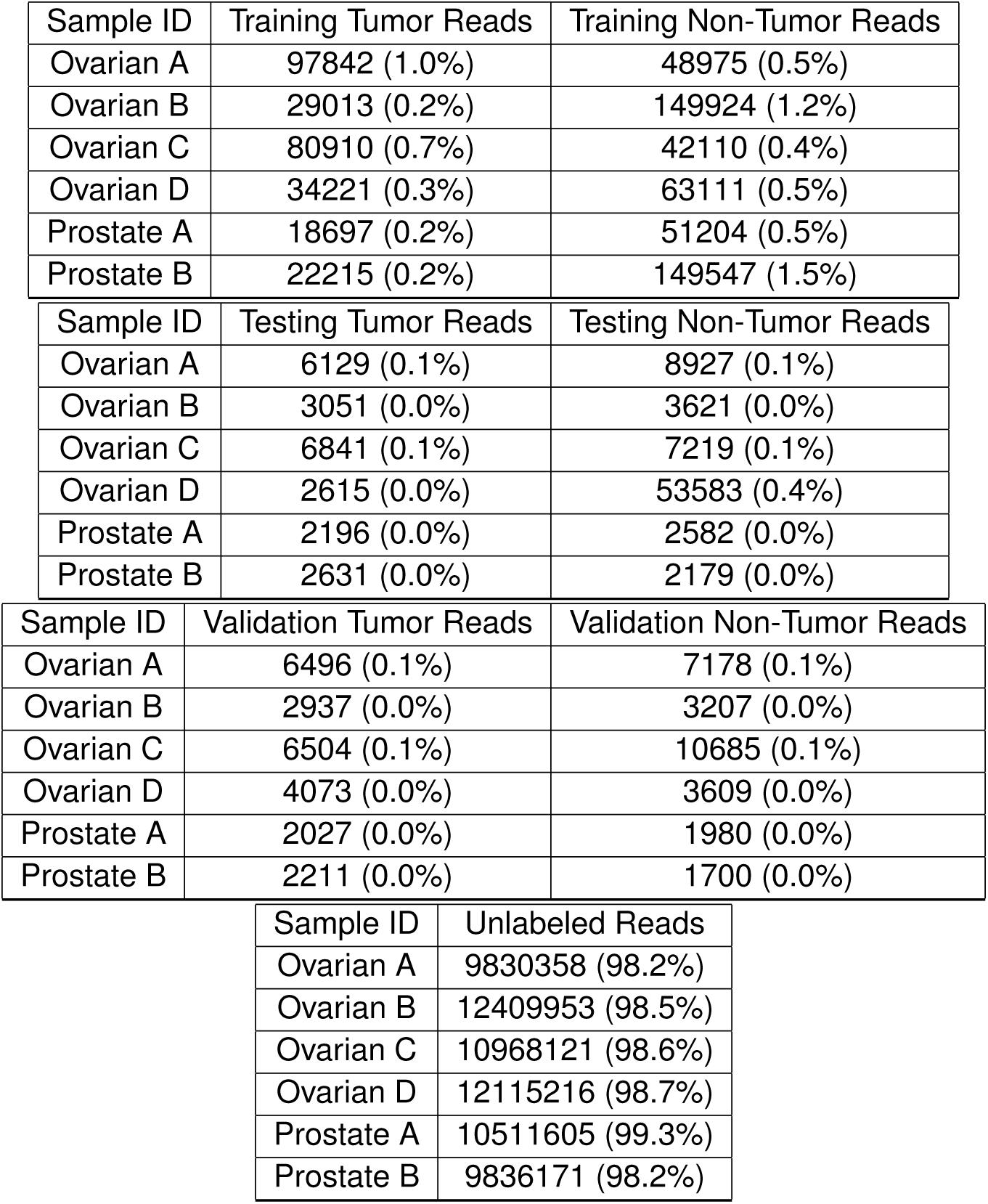
Classification counts for ground truth training data for the five samples in our cohort. The remaining reads are the reads for each sample where no ground truth origin label could be obtained. Percentages are expressed as a fraction of the total bulk tumor reads for the sample.

| Sample ID | Training Tumor Reads | Training Non-Tumor Reads |
| --- | --- | --- |
| Ovarian A | 97842 (1.0%) | 48975 (0.5%) |
| Ovarian B | 29013 (0.2%) | 149924 (1.2%) |
| Ovarian C | 80910 (0.7%) | 42110 (0.4%) |
| Ovarian D | 34221 (0.3%) | 63111 (0.5%) |
| Prostate A | 18697 (0.2%) | 51204 (0.5%) |
| Prostate B | 22215 (0.2%) | 149547 (1.5%) |

| Sample ID | Testing Tumor Reads | Testing Non-Tumor Reads |
| --- | --- | --- |
| Ovarian A | 6129 (0.1%) | 8927 (0.1%) |
| Ovarian B | 3051 (0.0%) | 3621 (0.0%) |
| Ovarian C | 6841 (0.1%) | 7219 (0.1%) |
| Ovarian D | 2615 (0.0%) | 53583 (0.4%) |
| Prostate A | 2196 (0.0%) | 2582 (0.0%) |
| Prostate B | 2631 (0.0%) | 2179 (0.0%) |

| Sample ID | Validation Tumor Reads | Validation Non-Tumor Reads |
| --- | --- | --- |
| Ovarian A | 6496 (0.1%) | 7178 (0.1%) |
| Ovarian B | 2937 (0.0%) | 3207 (0.0%) |
| Ovarian C | 6504 (0.1%) | 10685 (0.1%) |
| Ovarian D | 4073 (0.0%) | 3609 (0.0%) |
| Prostate A | 2027 (0.0%) | 1980 (0.0%) |
| Prostate B | 2211 (0.0%) | 1700 (0.0%) |

Supplementary Table 1: Classification counts for ground truth training data for the five samples in our cohort.
| Sample ID | Unlabeled Reads |
| --- | --- |
| Ovarian A | 9830358 (98.2%) |
| Ovarian B | 12409953 (98.5%) |
| Ovarian C | 10968121 (98.6%) |
| Ovarian D | 12115216 (98.7%) |
| Prostate A | 10511605 (99.3%) |
| Prostate B | 9836171 (98.2%) |

**Supplementary Table 2:**
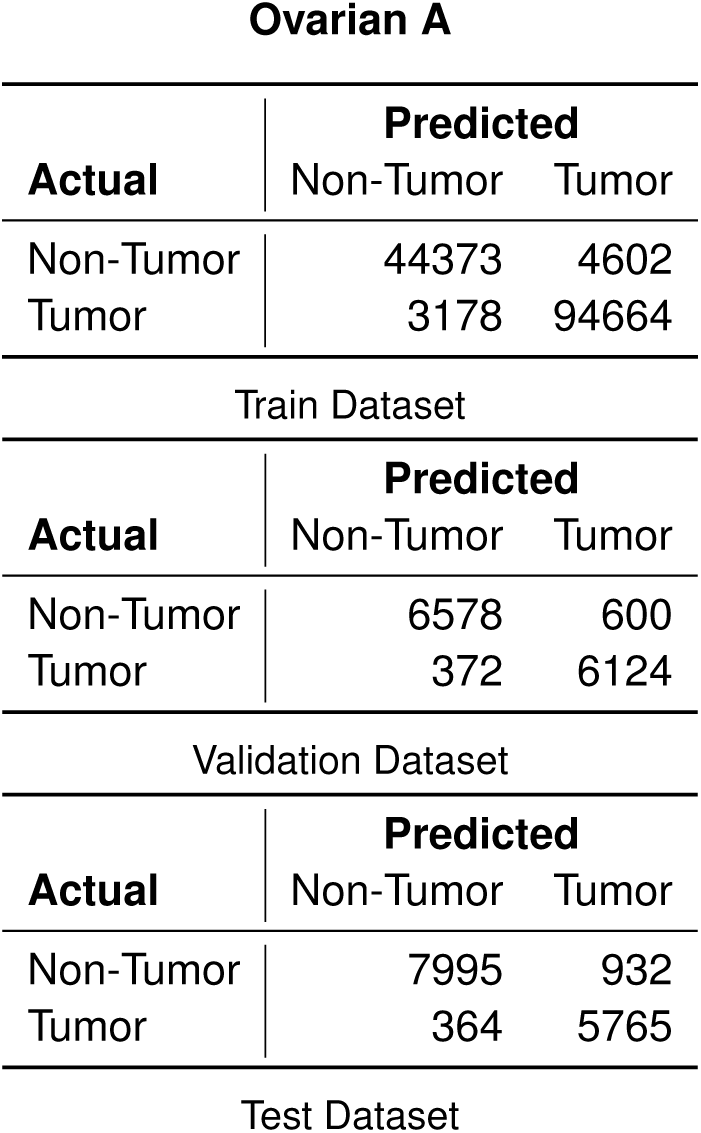
Confusion matrix for the labeled training, testing and validation datasets for the model trained and evaluated on sample ovarian A. A probability threshold of 0.5 was used for tumor read classification.

**Supplementary Table 3:**
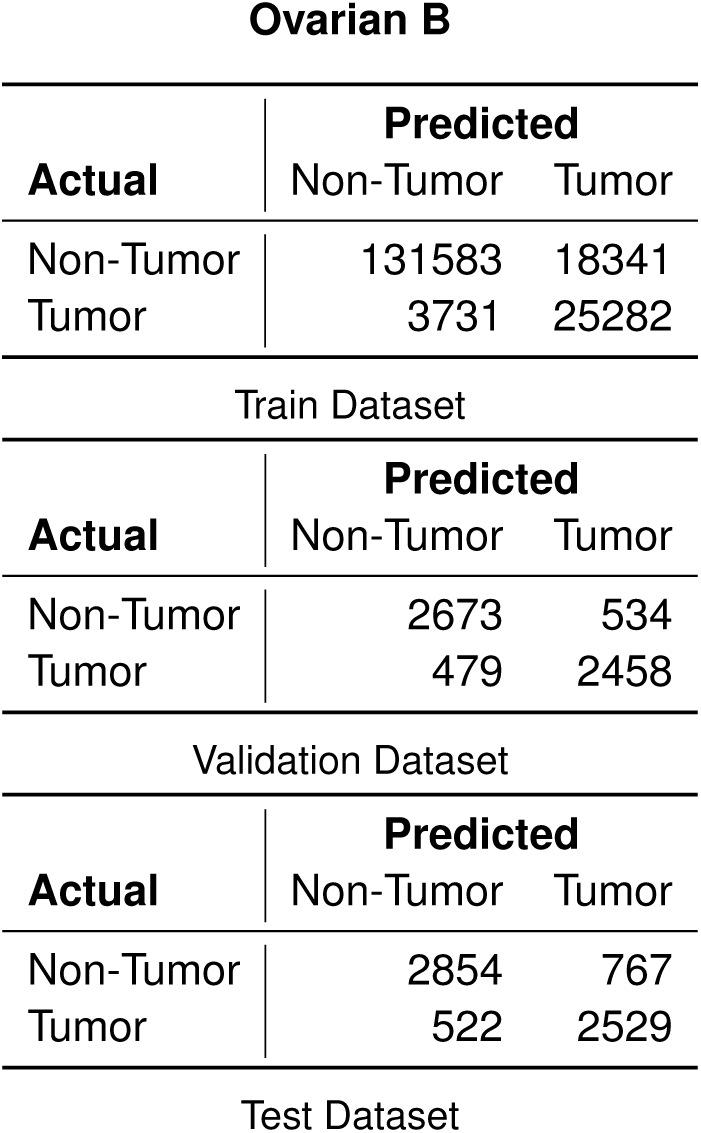
Confusion matrix for the labeled training, testing and validation datasets for the model trained and evaluated on sample ovarian B. A probability threshold of 0.5 was used for tumor read classification.

**Supplementary Table 4:**
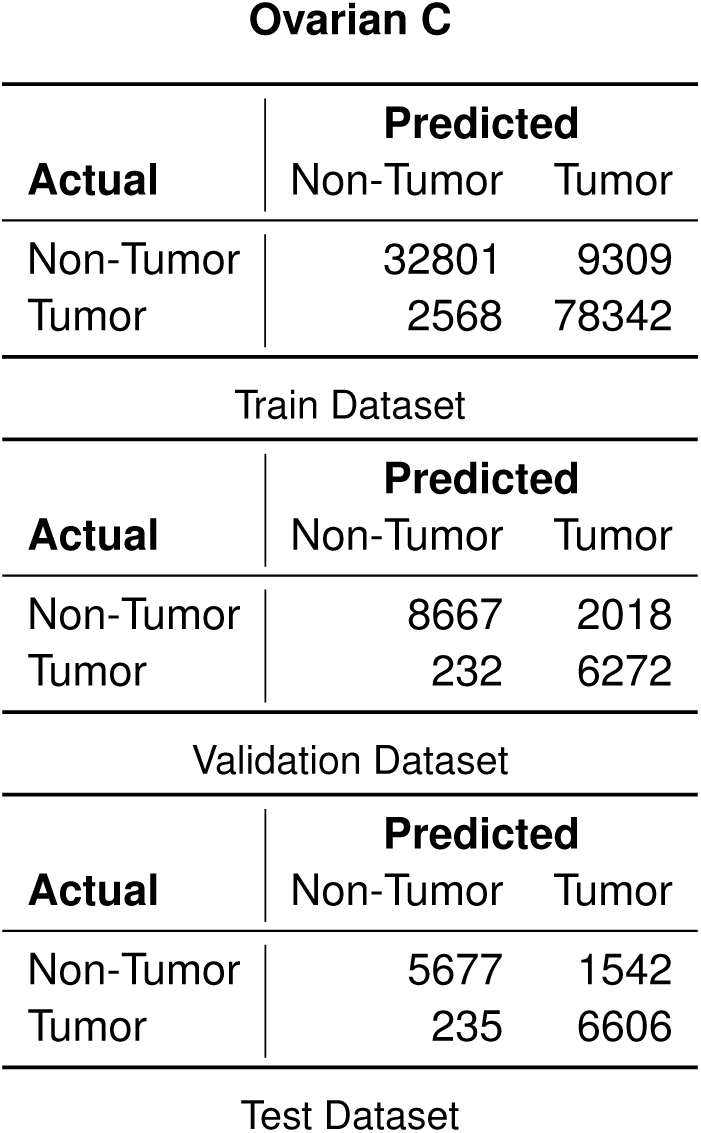
Confusion matrix for the labeled training, testing and validation datasets for the model trained and evaluated on sample ovarian C. A probability threshold of 0.5 was used for tumor read classification.

**Supplementary Table 5:**
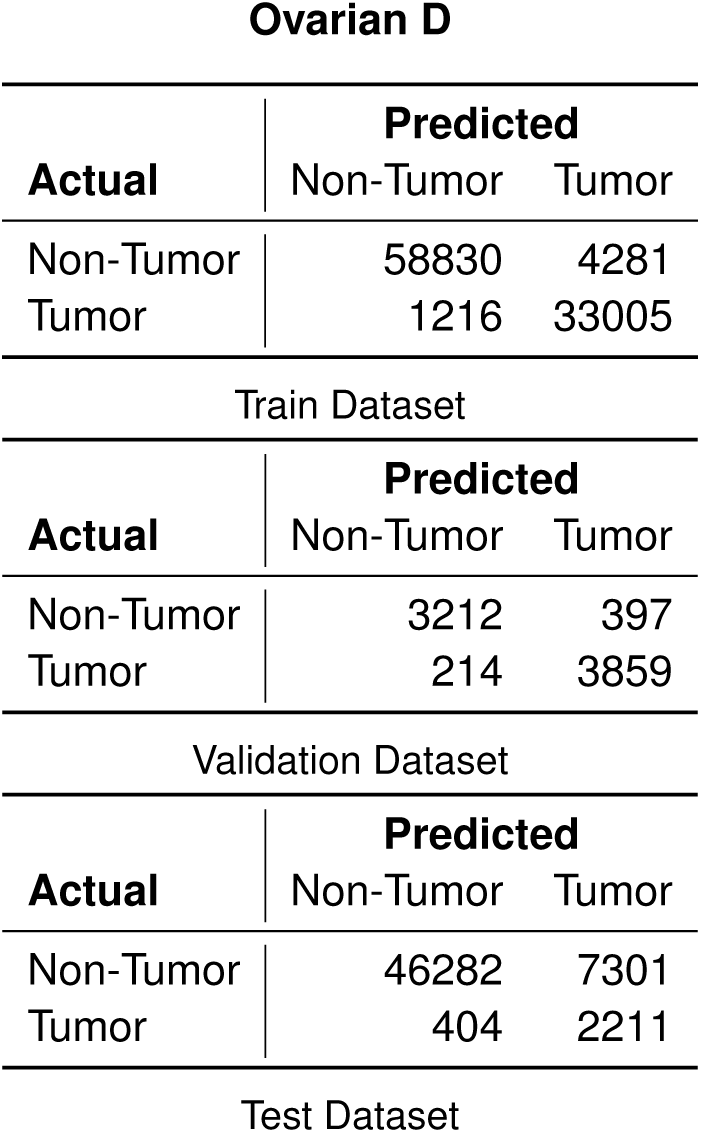
Confusion matrix for the labeled training, testing and validation datasets for the model trained and evaluated on sample ovarian D. A probability threshold of 0.5 was used for tumor read classification.

**Supplementary Table 6:**
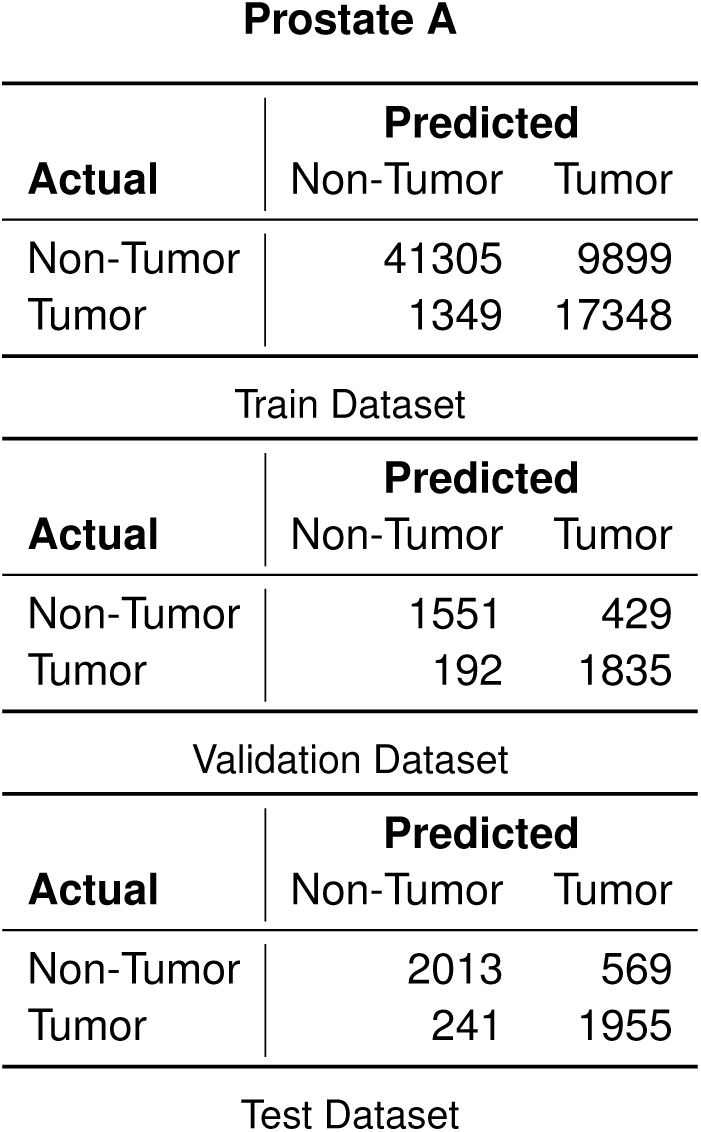
Confusion matrix for the labeled training, testing and validation datasets for the model trained and evaluated on sample prostate A. A probability threshold of 0.5 was used for tumor read classification.

**Supplementary Table 7:**
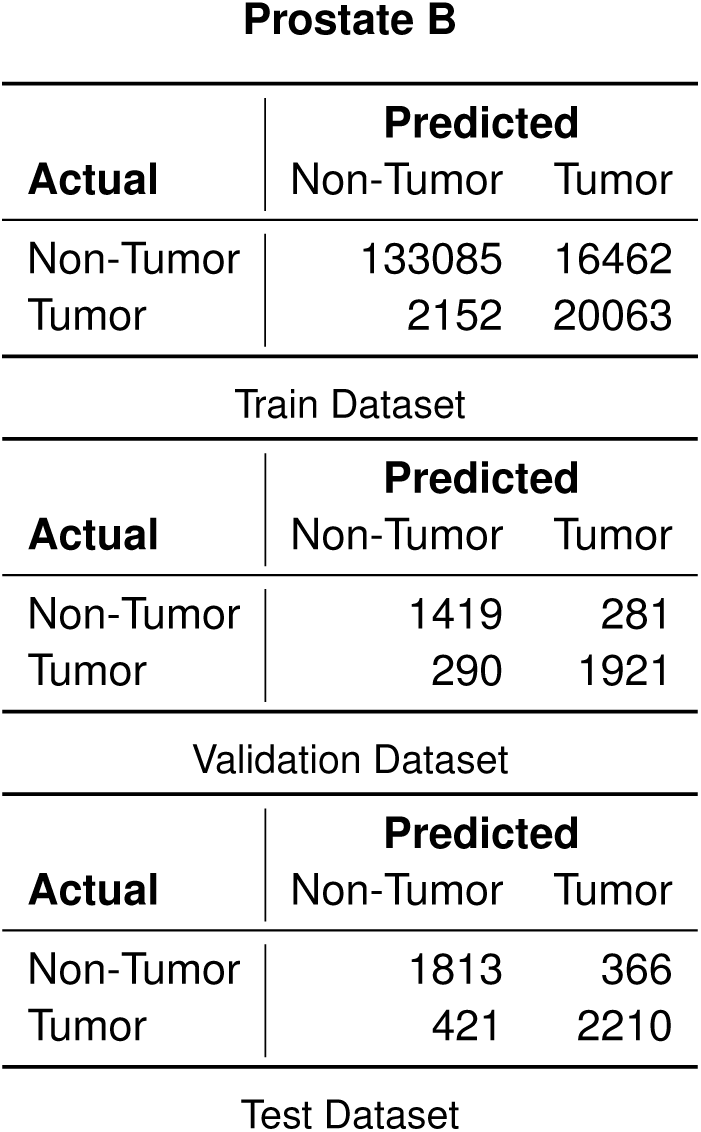
Confusion matrix for the labeled training, testing and validation datasets for the model trained and evaluated on sample prostate B. A probability threshold of 0.5 was used for tumor read classification.

